# Joint Vector Flow Mapping and Segmentation: Ill-Posedness, Differentiable Bayesian Inference, and Synthetic Vortex-Flow Benchmarks

**DOI:** 10.64898/2026.09.24.754220

**Authors:** Juan C. del Alamo, Cathleen M. Nguyen, Manuel Guerrero-Hurtado, Adithan Kandasamy, Bahetihazi Maidu, Andrew M. Kahn, Javier Bermejo, Pablo Martinez-Legazpi

## Abstract

Vector flow mapping (VFM) reconstructs left-ventricular (LV) blood velocity from color-Doppler echocardiography by combining the measured beamwise component with physical and regularizing constraints. Analysis of the discrete VFM formulation shows that the inverse problem is intrinsically ill posed: the occurrence of singular modes can be predicted from the geometry of the segmented blood-pool domain, the imposed boundary conditions, and the degree of smoothing. These modes can propagate uncertainty along entire transverse bands of the reconstructed velocity field, yet conventional VFM neither quantifies this uncertainty nor allows for correcting the blood-pool segmentation.

We introduce Bayesian VFM (B–VFM), a hierarchical framework that jointly infers radial and transverse velocities, a probabilistic blood-pool mask, their spatially resolved uncertainties, and hyperparameters weighting Doppler and segmentation fidelity, mass conservation, boundary conditions, and smoothness. The discretized posterior admits a closed-form gradient and exact Hessian, enabling computationally efficient, gradient-based MAP estimation, matrix-free Laplace sampling, and direct analysis of ill-posed modes. Posterior inference combines Gibbs sampling of conjugate Gamma-distributed hyperparameters with conditional maximum-a-posteriori estimation and a Laplace approximation for the high-dimensional velocity and mask fields. To accommodate systematic departures from planar mass conservation, B-VFM can learn the covariance of the planar divergence residual from an ensemble of flows and incorporate it as a structured model-discrepancy prior.

Independent chains converged reproducibly, while covariance priors learned from flow ensembles illustrated how model discrepancies can be incorporated into the inference. B–VFM was evaluated using Lamb–Chaplygin dipoles under ideal conditions and with Doppler corruption, Doppler voids, and segmentation defects, and using the Hicks–Moffatt family of spherical vortices to assess violations of planar mass conservation. The method produced smooth reconstructions, localized uncertainty near unreliable measurements and regions of model inconsistency, and used flow information to correct segmentation errors. Within the tested vortex family, the data-informed planar divergence prior reduced velocity bias and mask distortion. B–VFM thus provides an uncertainty-aware reconstruction method and a flexible foundation for future VFM formulations incorporating additional priors, observations, and physical models. Future work will evaluate the method using clinical data and more complex three-dimensional benchmark flows.

## 1 Introduction

In the past two decades, advances in medical imaging have allowed for enhanced visualization of the cardiovascular system. As a result, methods to measure intracardiac blood flow and analyze flow patterns have also improved [10, 31, 47]. Among the existing cardiovascular imaging modalities, echocardiography offers non-ionizing, inexpensive, portable imaging with short acquisition times. In particular, color-Doppler echocardiography remains the workhorse of clinical left ventricular (LV) flow evaluation. However, while it can be measured in linear, planar, or volumetric sectors, color-Doppler only senses the flow velocity in the direction of the ultrasound beam. Therefore, significant efforts have been devoted to inferring velocities in the cross-beam directions [34]. One of such techniques that is gaining widespread acceptance is vector flow mapping (VFM) [4, 21, 22, 33, 49, 52, 69, 76].

VFM processes color-Doppler data and a time-dependent segmentation of the LV wall in the apical long-axis view, also known as the three-chamber view. This echocardiographic view displays the mitral annulus, lateral LV wall, the apex, the LV outflow tract, and the aortic valve, making it ideal for observing the whole transit of blood through the LV. Each heartbeat, the dominant LV flow pattern typically alternates between a diastolic inflow jet flanked by a vortex ring, a telediastolic prograde swirling cell that redirects incoming blood towards the LV outflow tract, and a systolic jet [10]. Realizing that these features can be observed in the apical long-axis view [59], and that their dynamics were approximately planar [72] paved the way for VFM. Investigators have validated VFM on this plane using ground-truth data from CFD [4, 5, 33, 49, 74, 76] and *in vitro* models [2, 22]. Head-to-head validation of echocardiographic VFM measurements vs. phase-contrast MRI measurements has also shown favorable agreement in flow velocities in the left and right ventricles [9, 50]. Moreover, numerical descriptors of LV transport computed on the 3-chamber view correspond well with equivalent metrics computed from 4D flow MRI [54]. In addition to the LV, VFM has been evaluated in other vascular geometries, showing encouraging agreement with reference data from particle image velocimetry [3].

Early implementations of VFM estimated the cross-beam velocity by directly enforcing mass conservation along circular arcs of the color-Doppler sector together with no-penetration conditions at the LV walls [22, 33]. Subsequent developments combined these constraints with Tikhonov regularization, casting VFM as a least-squares estimation problem [4]. This least-squares formulation is equivalent to assuming that the residuals of each constraint follow a Gaussian distribution (i.e., a prior) with zero mean and uniform standard deviation, so that inference corresponds to maximum a posteriori (MAP) estimation [11]. The inverse of each prior’s standard deviation is a precision hyperparameter that acts as weight for the associated constraint. The number of hyperparameters can be reduced by introducing Lagrange multipliers to strictly enforce the physical constraints [76]. More recently, this method has been generalized to 3D inference from multi-plane Doppler acquisitions [75]. Several groups have proposed VFM reconstructions based on the streamfunction vorticity formulation, which is applicable in planar, incompressible flow [49, 52]. These methods have the convenience of implicitly introducing regularization by numerically solving a Poisson equation to determine the streamfunction although it may be challenging to extend them to 3D. Recent VFM methods also incorporate Doppler phase unwrapping [80].

Secondary analyses of VFM velocity fields can quantify LV flow transport topology [29], blood stasis [65], and fluctuating pressure fields [71]. Among other applications, VFM has been used to quantify the changes in LV flow caused by age [8], adverse LV remodeling [9, 13, 45, 57], pacemaker or left ventricular assistance device implantation and programming [62–64], and cardioembolic stroke risk [44, 60, 61]. Apart from the LV, several studies have applied VFM to quantify flow in the right ventricle [1, 16, 50], the aortic root [28, 39] and other regions of the cardiovascular system [30].

While VFM is an effective modality for quantifying LV flow patterns, most implementations enforce in-plane mass conservation and boundary conditions strictly, neglecting out-of-plane fluxes and errors associated with LV wall tracking. Least-squares implementations offer the possibility of balancing the residual from various physical constraints and regularization penalties [4] via the hyperparameter values that control each constraint’s weight. However, while heuristic approaches such as the L-curve criterion can inform the choice of hyperparameters [27], they add significant computational cost, lack robustness, and introduce arbitrariness in the solution. Adding to the lack of schemes to balance the different error sources, VFM uncertainty propagation is yet to be studied in detail. Naive VFM techniques considered error accumulation when integrating the mass conservation equation along circular arcs to choose weight functions for the integration [22] or to estimate posterior cross-beam velocity errors [70]. Nevertheless, these methods neglected the errors associated with wall tracking and regularization, and lumped all other errors together without distinguishing their origin.

Bayesian methods have emerged as a powerful tool for solving inverse problems while incorporating uncertainty quantification. For example, Sun et al. applied a Bayesian framework to optical flow estimation, providing spatially resolved confidence in motion fields [68]. Similarly, Borggaard et al. used Bayesian inference to estimate background flows from passive scalar data, highlighting the flexibility of probabilistic approaches in fluid estimation tasks [12]. More recently, Kontogiannis et al. [37] developed a Bayesian inverse Navier–Stokes framework to regularize 3D velocity fields and learn boundary position and parameters such as the viscosity coefficient.

This manuscript introduces B-VFM, a hierarchical Bayesian framework that jointly infers the velocity field, flow-domain mask, model weights, and their uncertainties. Because the B-VFM posterior is expressed entirely in discrete operator form, its gradient and Hessian can be derived exactly with respect to the latent variables. This differentiable structure supports efficient MAP optimization and curvature-based uncertainty propagation, while exposing the null modes and mask–velocity couplings governing VFM regularity. B-VFM also permits physical-model discrepancies to be learned from flow ensembles via a learned spatial covariance of departures from planar mass conservation from related three-dimensional flows. We evaluate the framework using controlled LC-dipole benchmarks with data and segmentation errors, followed by Hicks–Moffatt vortices to examine 3D model inconsistency and data-driven mass-conservation priors.

## 2 Methods

Bayesian VFM (B-VFM) combines physics-informed priors and input data uncertainty to infer LV flow velocity fields and correct the LV mask. It extends recent least-squares approaches that yield a single best-fit solution by minimizing a cost function [4, 5, 76] by treating the velocity 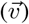 and LV mask (*m*) as random variables. B-VFM relies on a probability model that encodes our beliefs about the likelihood of different solutions, and sample the posterior distribution conditioned on the observed data. The final estimates correspond to the mean of the posterior distribution of 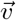 and *m*, and their uncertainty estimates correspond to the posterior’s standard deviation.

### 2.1 Bayesian Formulation of Vector Flow Mapping

#### 2.1.1 Effect of Blood-Domain Geometry and Priors on VFM Ill-Posedness

Adopting a cylindrical coordinate system with its origin at the ultrasound transducer (Figure 1) B-VFM can be formulated as the inversion of a linear transformation relating the measured color-Doppler velocity (*V*) and LV mask (*M*) to the true radial and transverse velocities (*v*_*r*_ and *v*_*θ*_), and the LV mask, i.e.,

**Figure 1:**
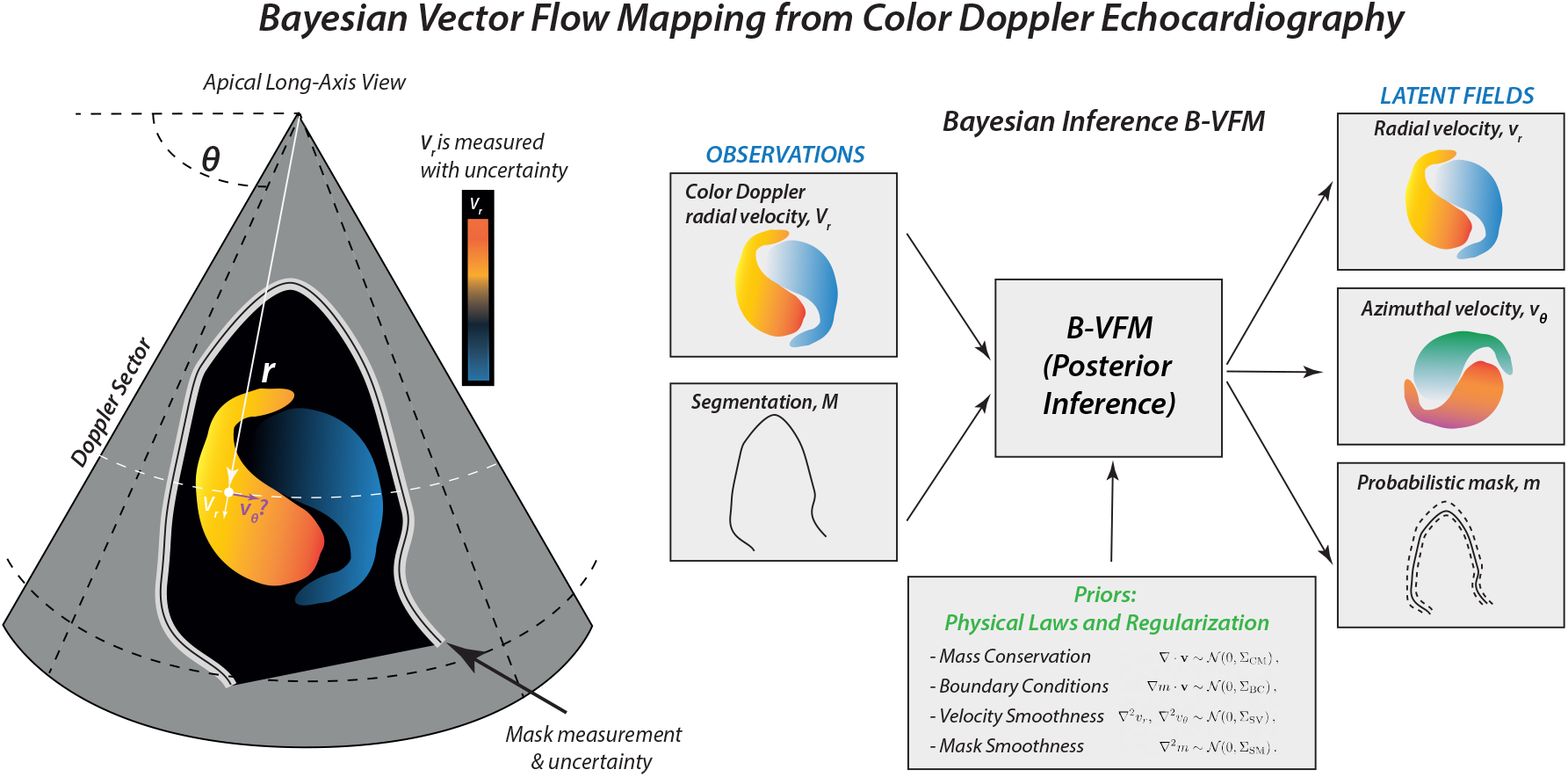
Schematic showing an overview of the Bayesian Vector Flow Mapping (B-VFM) method.

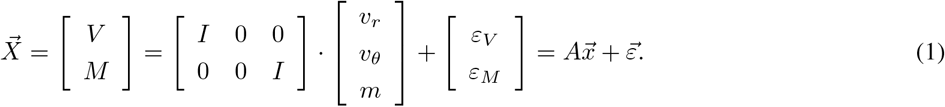

Discretizing this transformation on a regular mesh with *N*_*r*_ and *N*_*θ*_ points in the radial and angular directions, the 3*N*_*r*_*N*_*θ*_ *×* 1 parameter vector,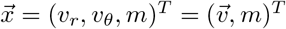, contains the values of the velocity components and LV mask indicator function at each pixel of the image. The 2*N*_*r*_*N*_*θ*_ *×* 1 measurement vector is 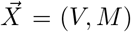, where *V* contains color-Doppler velocity values and *M* indicates whether a pixel belongs to the flow domain Ω. In a zero-uncertainty scenario, *M* is a binary mask containing ones and zeros whose boundary, ∂Ω, delineates, e.g., the LV endocardium. More broadly, *M* is a probabilistic atlas obtained from a segmentation algorithm [18] or by averaging several binary masks delineated by different users or methods. The measurement errors vectors *ε*_*V*_ and *ε*_*M*_ are assumed to be white noise, so that *V* and *M* follow normal distributions with zero mean and diagonal covariance matrices Σ_*V*_ and Σ_*M*_,

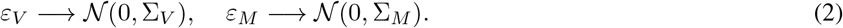

These two matrices can be considered as known if the Doppler and segmentation errors are available from the imaging pipeline. Alternatively, they can be considered hyperparameters in the hierarchical Bayesian approach (see §2.2). In the latter case, it is convenient to model 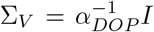 and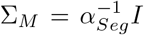, where *α*_*DOP*_ and *α*_*Seg*_ are the corresponding precision hyperparamters. This model assumes spatially homogeneous white noise, but more involved spatial models considering, e.g., the depth-dependence of Doppler measurements, or spatial correlations feeding the off-diagonal terms of Σ_*V*_ and Σ_*M*_, would also be possible. In particular, it is reasonable to assume that *α*_*Seg*_ is high in regions well inside and well outside the LV mask regardless of the segmentation method.

The linear transformation in equation (1) is represented by a rectangular matrix, *A*, whose *I* and 0 blocks are respectively identity and square null matrices of size *N*_*r*_*N*_*θ*_. Alternatively, the *I* blocks could be replaced by discrete interpolant or convolution operators if one wishes to infer 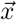 on a different mesh than that of 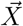. In any case, because *A* is rectangular and *A*^*T*^ *A* is singular, the transformation cannot be inverted even in least squares sense without injecting prior information about *v*_*θ*_ and the other two variables. A classic VFM prior is to consider that the divergence of the velocity field in Ω follows a normal distribution of zero mean and diagonal covariance matrix

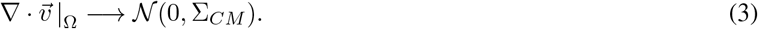

In discrete form, this prior can be written as

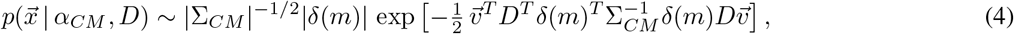

where 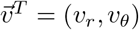, *D* is a *N*_*r*_*N*_*θ*_ × 2*N*_*r*_*N*_*θ*_ matrix representing the discrete divergence operator in cylindrical coordinates, and *δ*(*m*) is a diagonal matrix of size *N*_*r*_*N*_*θ*_ that indicates the pixels belonging to the blood pool, Ω. The divergence covariance matrix for this prior is modeled as 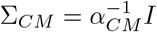where *α*_*CM*_ is a precision hyperparameter. This model assumes that departures from zero divergence are spatially homogeneous and uncorrelated. Empirical knowledge of the spatial structure of 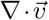 from, e.g., phase contrast MRI or computational fluid dynamics, could be used to further inform this model (see §2.3.1). Here, the complete representation of the divergence operator including the *r*^*−*1^ metric, i.e., 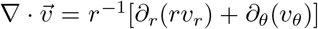, should be used to prevent the appearance of *r*-dependent errors. Mathematically, it is straightforward to see that not including the metric term is equivalent to assuming that *α*_*CM*_ *∼ r*^2^. Previous VFM methods often neglect the *r*^*−*1^ metric by multiplying both sides of the 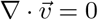 equation by *r*, but this simplification lacks justification unless zero divergence is strictly enforced.

Additional priors include wall boundary conditions for mass conservation and smoothness constraints on the velocity field and the LV wall. Wall boundary conditions for mass conservation imply that blood velocity must be parallel to the segmented LV wall, i.e., 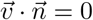 where 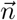 is the vector normal to ∂Ω. Thus, we consider

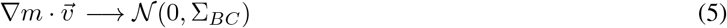

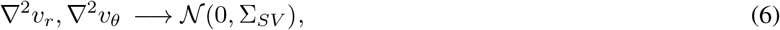

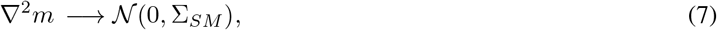

where the covariance matrices 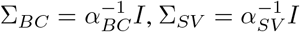, and 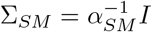 are determined by their respective precision hyperparameters, *α*_*BC*_, *α*_*SV*_, and *α*_*SM*_. The corresponding discretized probability functions are:

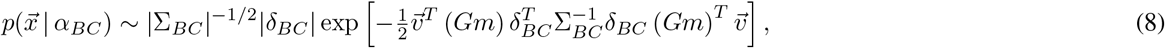

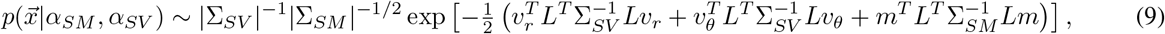

where *L* = *DG* is the discretized Laplacian operator, and *δ*_*BC*_ is a diagonal indicator matrix that zeroes out pixels near open boundaries of the flow domain, such as cardiac valves. This matrix can also be tailored to exclude enforcement of 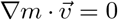 in regions known *a priori* to lie far from the LV endocardium, septal defects, LVAD inlet cannulas, etc.

Therefore, we model the final prior distribution including all physical and smoothness constraints as

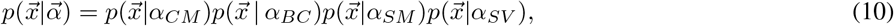

and, assuming independent Doppler and mask errors, the likelihood of our observations as

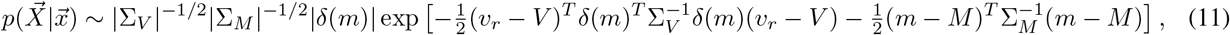

where notation has been compacted using the hyperparameter vector 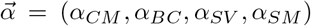, measurement noise vector 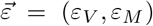, and operator vector *O* = {*D, G, δ*_*BC*_}. As stated above, since a reliable estimate of 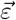 may not be always available, we will generally consider it to be inferrable, expanding the hyperparameter vector to 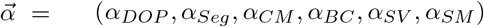. In section §2.2, this framework is used to find maximum a posteriori (MAP) estimates of 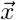 that maximize the conditional posterior 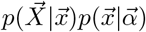 which, together with marginalization over 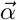, allows us to infer expected values and variances of 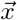. For simplicity and generality, we assume the Doppler map *V* has already been unwrapped. Appendix §SI3 provides probability models for phase unwrapping, showing phase unwrapping can be performed as a standalone preprocessing step prior to running B-VFM while retaining Bayesian rigor.

### 2.2 Hierarchical Bayesian Framework

We adopt a hierarchical Bayesian formulation to infer the unknown field 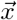 along with hyperparameters 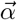 that modulate the strength of the prior constraints. In this framework, the hyperparameters are themselves assigned hyperpriors 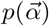 to reflect uncertainty, and our goal is to estimate the posterior marginals 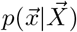 and 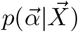, or at least compute their expectation and variance by integrating the joint posterior distribution,

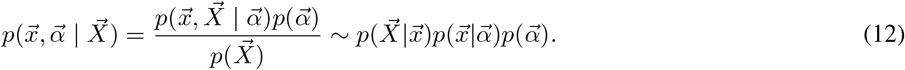

The joint posterior cannot be evaluated directly due to the coupling between 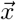 and 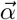, and the intractability of the marginal likelihood 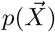. However, the models developed in §2.1 enable the construction of Markov Chain Monte Carlo (MCMC) methods that iteratively sample from known conditional distributions, creating chains that converge to the joint posterior [23]. Each MCMC chain is initialized with random hyperparameter values. If needed to promote convergence, plausible starting values can instead be estimated from a velocity field reconstructed using a standard VFM method, such as *i*VFM [4], as described in §2.2.1 below. To ensure robustness, an initial segment of each chain (burn-in) is discarded prior to inferring expectation and variance. We also run multiple MCMC chains with different hyperparameter initializations and evaluate whether algorithm consistently converges to similar posterior distributions.

#### 2.2.1 MAP-Based Laplace-Approximate Gibbs Sampler

The Gibbs sampler is a well-known method to create MCMC chains that converge to the joint posterior [14, 58]. In the iteration *it*, 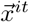 is sampled by drawing from 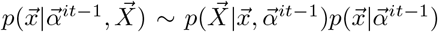, and 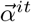 from 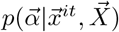. If the hyperprior 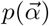 is chosen carefully (e.g., a Gamma distribution), then the hyperparameter conditional posterior 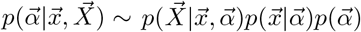 retains the same functional form as 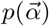 and can be obtained in closed form. This property is widely known as conjugacy [23].

Direct sampling from 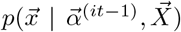 is not tractable in B-VFM because the residuals entering several priors—most notably the boundary-condition prior in Eq. (8)—depend nonlinearly on 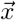, rendering the conditional posterior non-Gaussian.

At each iteration, we therefore construct a Laplace approximation centered at the conditional maximum a posteriori (MAP) estimate, 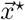, as summarized in Algorithm 1, and draw 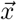 from the resulting Gaussian distribution. For fixed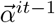, we first minimize the negative conditional log-posterior,

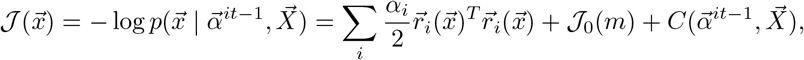

where 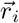 is the residual of the *i*-th constraint, *J*_0_(*m*) arises from the mask-dependent normalization factors in equations (8-11), and *C* is a conditional constant (see §SI1 in the Appendix for a full derivation). Minimization yields the conditional MAP estimate 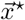, from which a posterior sample 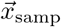 is drawn from the Gaussian distribution 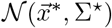 Here, 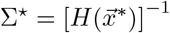 and 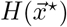 is the Hessian of the negative conditional log-posterior evaluated at the MAP estimate, which represents the local posterior precision. A closed-form expression for the precision matrix *H* is provided in §3.1.1. While, in principle, this expression should permit obtaining Σ^*⋆*^, its large size 3*N*_*r*_*N*_*θ*_ *×* 3*N*_*r*_*N*_*θ*_ makes direct matrix inversion impractical.

Therefore, we work with the Gauss–Newton approximation

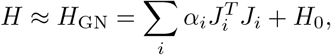

where 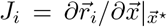 is the residual Jacobian and *H*_0_ is a positive diagonal matrix representing the local curvature of *J*_0_(*m*). We present analytical expressions for the exact and Gauss–Newton Hessians in the Appendix (§ SI2), showing that they differ only through residual-dependent corrections to the velocity–mask cross-blocks. Thus, the error introduced by the Gauss–Newton approximation is expected to be smallest near the MAP estimate and is confined to the coupling between the inferred velocity and mask.

Next, a normally distributed perturbation ***η*** is generated using independent vectors 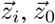 drawn from *N*(0, *I*),

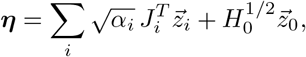

which ensures that Cov(***η***) = *H*_*GN*_. Then, a perturbation 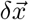 is obtained by solving 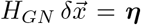 and a conditional field sample can be built from 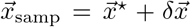 where Cov 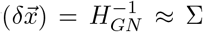. Thus, a normally distributed sample can be generated without assembling, inverting, or factorizing *H*. Additionally, we accelerate the linear solve using a preconditioned conjugate gradient method with the Jacobi preconditioner diag(*H*_*GN*_).

Hyperparameter values are subsequently sampled using Gamma distributions for hyperparameters, i.e., *p*(*α*) = Γ(*α*| *β, ϕ*) *∼ α*^*β−*1^*e*^*−αϕ*^. The Gamma hyperprior is convenient due to its conjugacy with the conditional priors, yielding conditional posteriors that are also Gamma distributions:

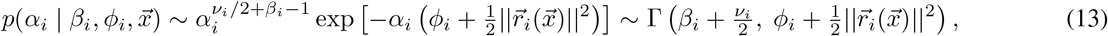

where *ν*_*i*_ = rank(*J*_*i*_) is the number of independent residual components for each constraint. The tractability of the Gamma posterior allows for direct sampling in MCMC.

In addition to enforcing positivity, the Gamma distribution offers a flexible means of encoding prior beliefs through its shape and rate parameters, *β*_*i*_ and *ϕ*_*i*_. For simplicity, we adopt non-informative hyperpriors with *β*_*i*_ = 1 and *ϕ*_*i*_ = 10^*−*5^. In the non-informative limit—i.e., when 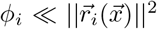 and *β*_*i*_ *≪ ν*_*i*_, the resulting posterior distribution for *α*_*i*_ has mean and variance given by

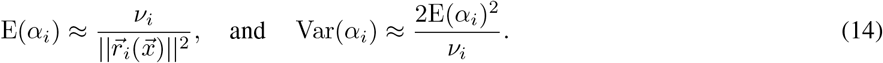

Because the rank *ν*_*i*_ *∼ N*_*r*_*N*_*θ*_ is large in B-VFM applications, the resulting conditional posteriors of *α*_*i*_ are sharply peaked and yield stable, data-adaptive regularization. This procedure tends to balance the contributions of the weighted terms, thereby inducing an implicit model selection mechanism: components associated with large residuals have their weights shrunk toward zero, while those that capture meaningful structure are retained [23].

##### Algorithm 1

MAP-Based Laplace-Approximate Gibbs Sampler

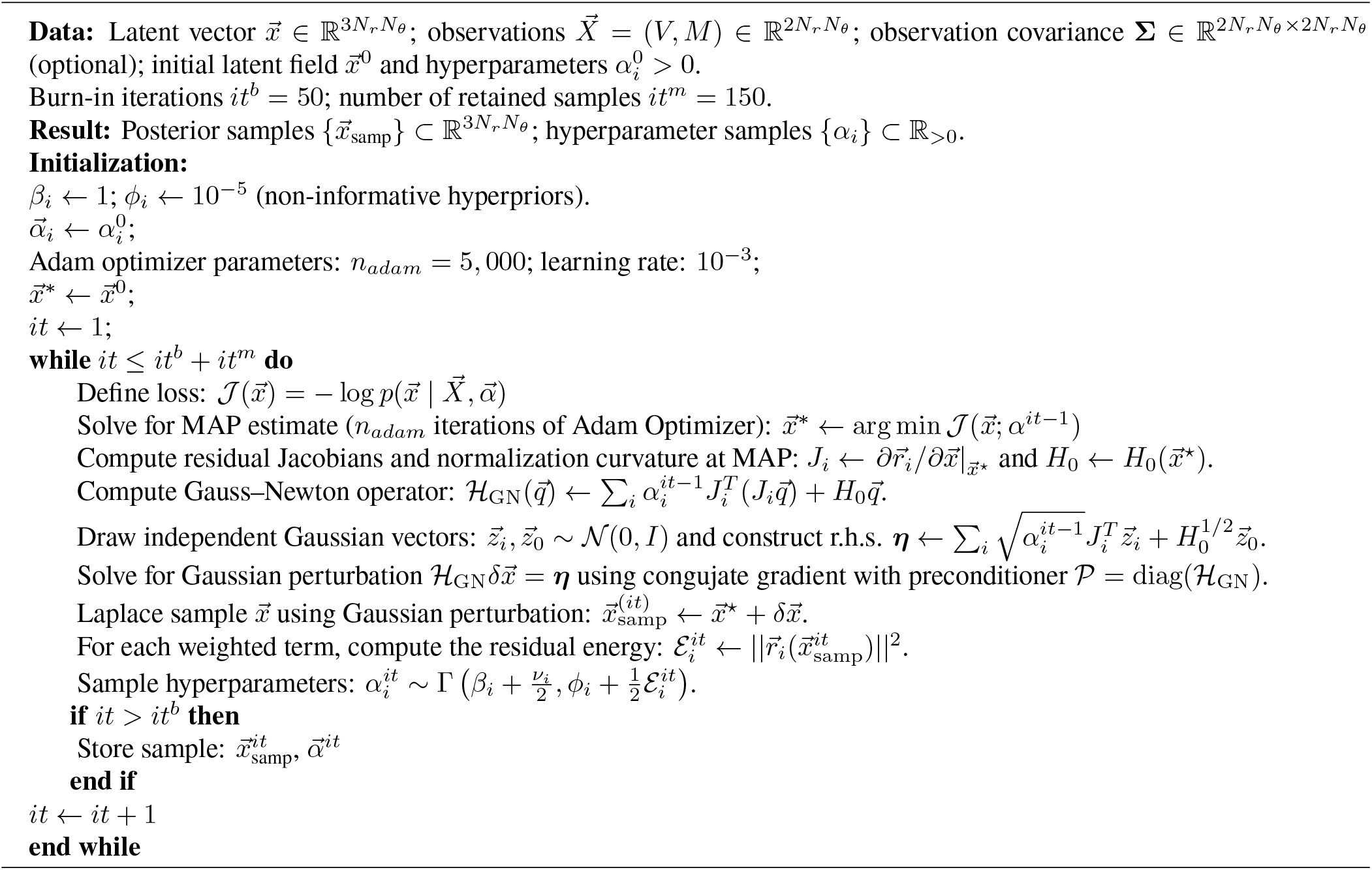

#### 2.2.2 Adam Optimizer

The implementation of Algorithm 1 requires calculating the conditional MAP estimate of 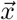 at each iteration. We obtained this estimate using Adaptive Moment Estimation (Adam) applied to the complete conditional negative log-posterior and its analytical gradient (analytically derived in §SI1). We used initial learning rates of 10^*−*2^ and 10^*−*3^ for the velocity and mask variables, respectively. Both were reduced using a standard inverse-time schedule at a relative rate of 10^*−*4^ per iteration. Standard values of momentum parameters were used for smoothing the gradient and squared gradient (*β*_1_ = 0.9 and *β*_2_ = 0.999, respectively). The optimization was stopped after 5,000 iterations; increasing this parameter did not seem to significantly affect B-VFM results.

#### 2.2.3 Optional Hard Mask Constraints

Our BVFM implementation optionally supports hard constraints at pixels whose segmentation label can be determined unambiguously, such as pixels located well inside or well outside the segmented boundary. When this option was used, the constraint regions were constructed from the segmented mask by applying dilation and erosion operations with a width equal to (20%) of the domain size. Pixels outside the dilated mask were designated as known-outside, whereas pixels inside the eroded mask were designated as known-inside. Pixels between the eroded and dilated boundaries were treated as uncertain. Mask values were fixed at (1) and (0) for known-inside and known-outside pixels, respectively, whereas the mask was inferred over the interval ([0,1]) at uncertain pixels. These constraints were imposed after every Adam update. The corresponding mask coordinates were also excluded from the Gaussian perturbation in the Laplace sampling step, ensuring that their prescribed values remained unchanged in every posterior sample.

### 2.3 Benchmark Vortex Flows

Vortex rings are a hallmark of LV diastolic flow, and abnormalities in their formation have been linked to LV dysfunction [20, 24, 45, 57]. Therefore, to evaluate B-VFM under controlled conditions, we used synthetic flows containing vortical structures representative of a planar cut through a vortex ring. Our primary benchmark was the Lamb–Chaplygin (LC) dipole [48], a translating solution of the 2D incompressible Euler equations consisting of a steadily propagating, counterrotating vortex pair. Because the LC dipole satisfies 2D mass conservation exactly, B-VFM is expected to recover *v*_*θ*_ and *m* up to numerical discretization error when supplied with error-free Doppler and mask data. This flow also provides a controlled benchmark for evaluating the effects of errors in these input fields.

To represent the LC dipole, we use a polar coordinate system with origin at the dipole center. To avoid confusion with the ultrasound probe based polar coordinates, we denote (*ρ, ϑ*) and (*u*_*ρ*_, *u*_*ϑ*_) the dipole-based coordinates and velocity components. In this reference frame, the streamfunction is given by

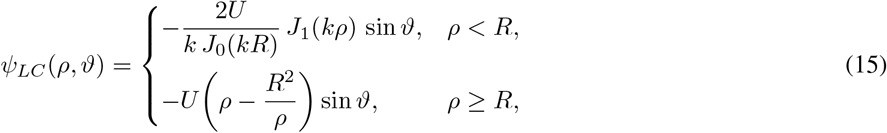

where *U* is the dipole translation speed, *R* is its radius, and *J*_*n*_ denotes the *n*th-order Bessel function of the first kind. The parameter *k* is selected such that *kR* = *α*_1_ ≈ 3.8317, where *α*_1_ is the first zero of *J*_1_. This choice ensures continuity of the streamfunction and velocity across *ρ* = *R*. The vorticity is confined to the dipole interior, while the exterior solution approaches the uniform far-field velocity associated with the translating reference frame.

To evaluate B-VFM under quantifiable departures from its planar-flow assumption, we also considered the Hicks–Moffatt (HM) family of swirling spherical vortices [51]. These are axisymmetric, 3D incompressible flows parameterized by the dimensionless radial wavenumber *κ*. Hill’s spherical vortex is recovered in the limit *κ* → 0, whereas nonzero values of *κ* modify the internal meridional flow and introduce an azimuthal velocity component.

In this case, the stream function is [51]

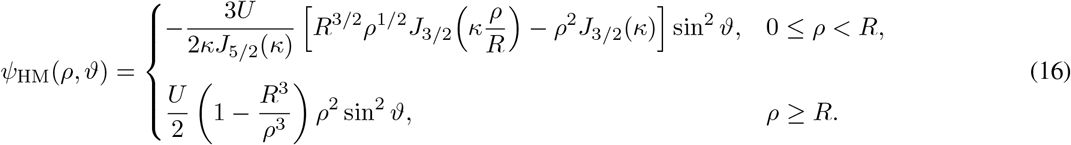

Changing the sign of *κ* simply reverses the swirl direction, so we consider only *κ* ≥ 0. Specifically, we used *κ* ∈ [0, 5.4], below first positive zero of *J*_5*/*2_, *κ*_1_ ≈ 5.7635. As *κ* is increased beyond this value, additional internal recirculating cells appear and the meridional flow no longer resembles a vortex pair.

In all simulations we used *R* = 5 cm and *U* = 5 cm/s. For the HM vortices, each 3D velocity field was sampled on a meridional plane containing the vortex symmetry axis and mapped onto a fan-shaped polar grid representative of an echocardiographic acquisition. The domain spanned depths of 0.4 cm ≤ *r* ≤ 12 cm and an angular sector of 106^*°*^. The vortex center was located at (*r, θ*) = (7 cm, 0). The in-plane radial component *v*_*r*_ was supplied to B-VFM as the input color-Doppler measurement, *V*, whereas the in-plane transverse component *v*_*θ*_ was withheld from the inference and retained as ground truth for validation. The ground-truth blood-domain mask was defined as *M* = 1 inside the dividing streamline *ρ* = *R* and *M* = 0 outside.

Although each HM vortex satisfies 3D mass conservation exactly, its meridional-plane velocity generally does not satisfy the 2D continuity equation assumed by conventional VFM. In the vortex-centered coordinates (*ρ, ϑ*), axisymmetric incompressibility implies

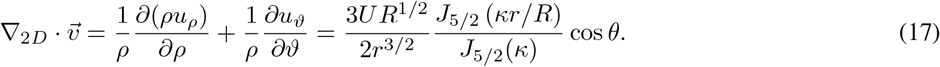

The left-hand side is the 2D divergence of the velocity observed in the meridional plane, whereas the right-hand side contains the geometric contributions absent from a planar formulation.

Figure 2 compares the LC flow field with two representative members of the HM family, corresponding to *κ* = 0 (Hill’s spherical vortex) and *κ* = 5. Although all three fields exhibit a vortex-pair structure, their velocity and vorticity distributions differ markedly. Moreover, whereas the LC dipole is exactly divergence-free in 2D, the planar divergence,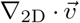, is nonzero for both HM vortices. As *κ* increases from 0 to 5, the divergence becomes more spatially concentrated and displaces farther from the vortex boundary.

**Figure 2:**
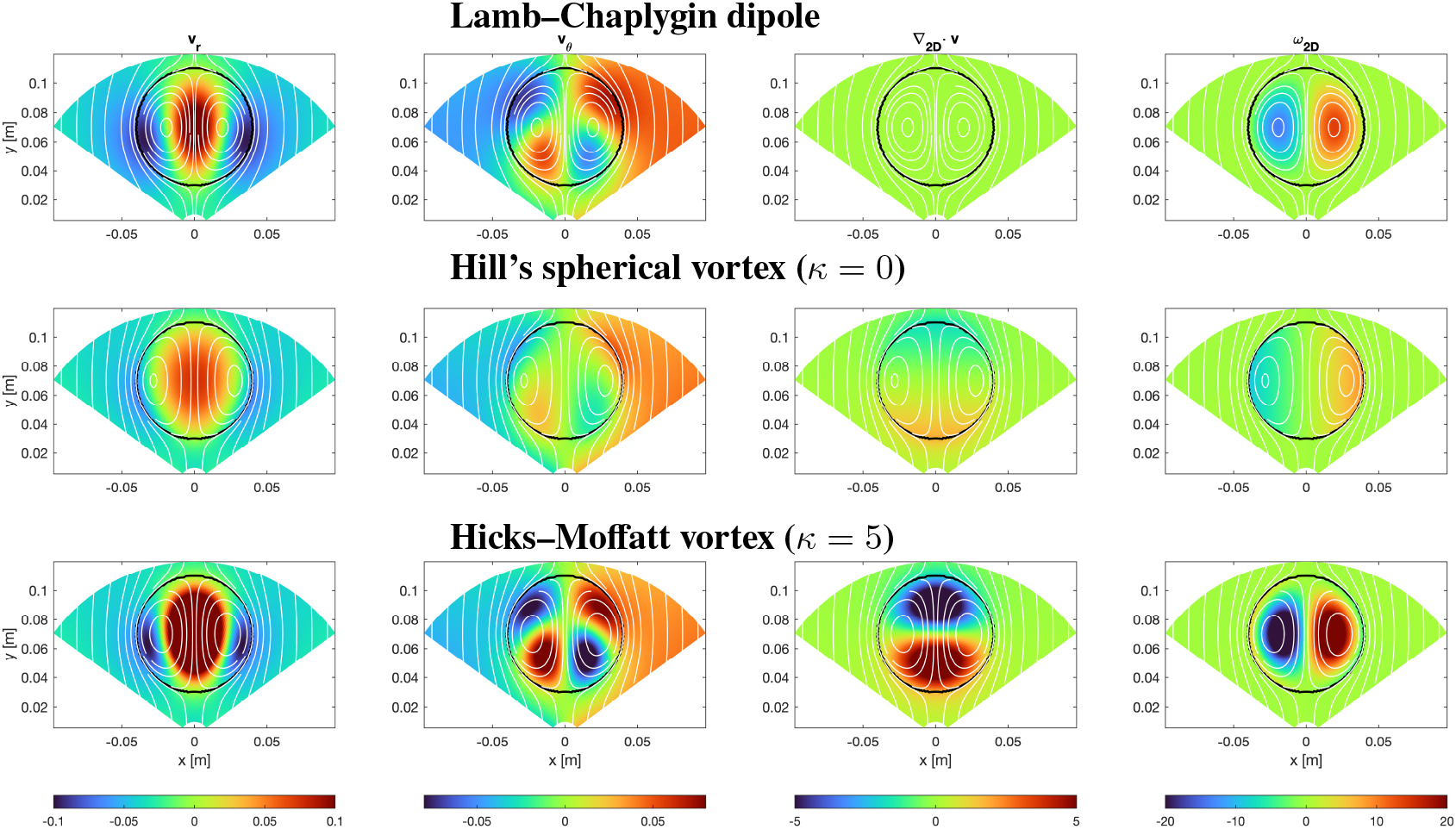
Representative synthetic vortex flows used to evaluate BVFM. Columns show, from left to right, the radial velocity (*v*_*r*_; m s^*−*1^), polar velocity (*v*_*θ*_; m s^*−*1^), planar divergence 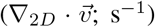, and out-of-plane vorticity (*ω*_2*D*_; s^*−*1^). Rows show, from top to bottom, the Lamb–Chaplygin two-dimensional dipole, Hill’s spherical vortex (the *κ* = 0 member of the Hicks–Moffatt family), and the Hicks–Moffatt vortex with *κ* = 5. In each panel, the black contour denotes the vortex boundary and the white curves denote streamlines.

**Figure 3:**
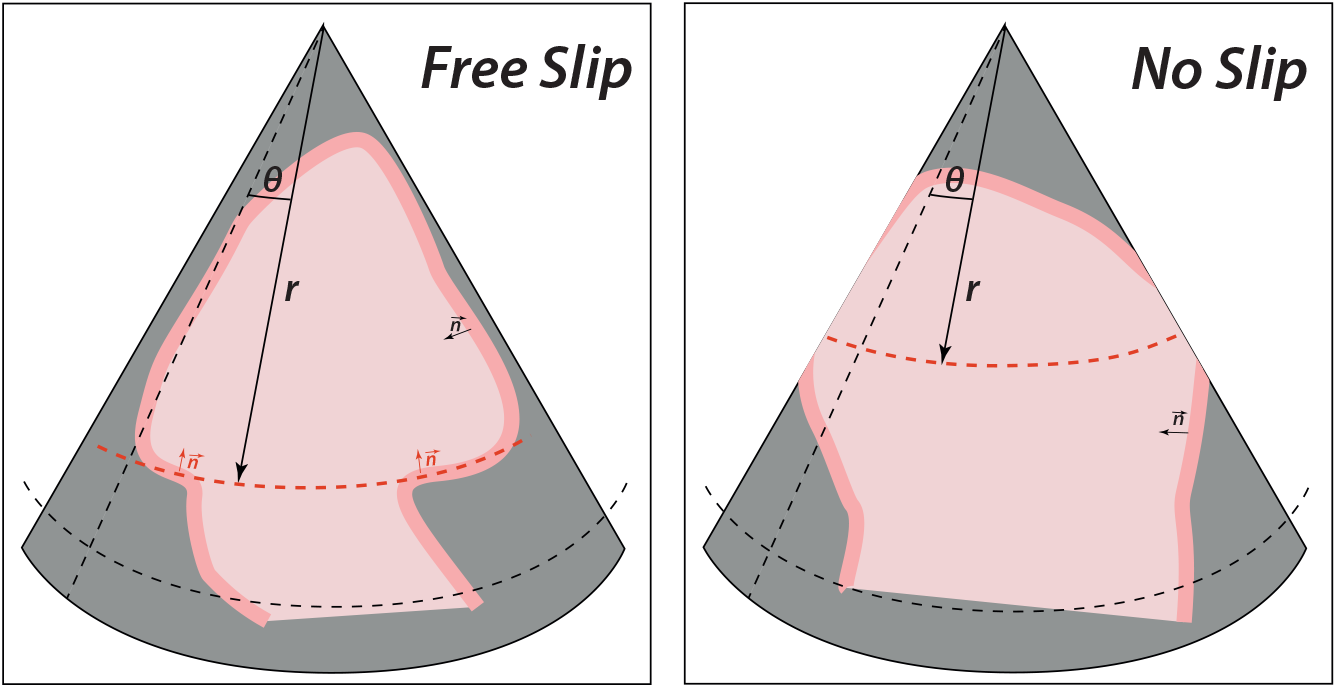
Schematic showing the scenarios in which VFM with mild or no within-inference velocity smoothing becomes prone to banded artifacts depending on type of boundary conditions and blood-pool mask geometry. Left: Under free-slip (no penetration) boundary conditions, VFM can display artifacts along constant-r arcs for which the mask boundary is completely aligned with the *θ*-direction (red dashed line). Right: Under no-slip boundary conditions, VFM can display banded artifacts along constant-r arcs with no boundary conditions.

#### 2.3.1 Computationally efficient data-informed prior for planar mass-conservation discrepancies

To incorporate non-planar flow information into B-VFM, we learned the statistics of 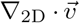 from an ensemble of HM vortices. Specifically, we generated *K* = 40 ensemble realizations with *κ* uniformly spaced between 0 and 5.4. To isolate changes in flow structure from changes in velocity magnitude, each realization was rescaled to have the same root-mean-square velocity as the *κ* = 0 vortex.

For each case, we computed 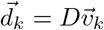, using the same discrete divergence operator *D* used for B-VFM and its ensemble mean, 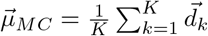, was defined as the expected divergence in the mass-conservation prior. We then formed the matrix *U* whose *k*-th column is the mean-centered divergence field, 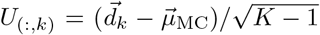, so that *UU*^*T*^ is the empirical covariance of the divergence fields. To regularize this covariance, we applied a small amount of shrinking toward its diagonal and added diagonal loading, yielding

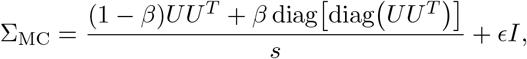

where *s* is the mean diagonal of *UU*^*T*^, and *β* = *ϵ* = 0.1. The resulting prior was

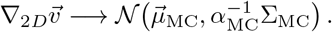

As explained above, direct sampling requires inverting this dense covariance matrix every Gibbs iteration, which is prohibitively expensive. Moreover, one cannot generally rely on having an analytical expression for an empirically derived Σ_MC_. However, one can exploit the particular structure of the covariance,

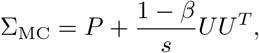

where

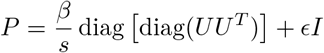

is a diagonal matrix and *U* has low rank, i.e., the number of realizations *K* used for learning. Defining *W* = *P* ^*−*1*/*2^*U*, we applied the Woodbury identity,

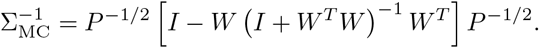

Thus, only a *K × K* matrix is inverted, avoiding storage and inversion of the full spatial covariance.

## 3 Results

### 3.1 Ill-posedness and Regularization of VFM

Vector Flow Mapping (VFM) poses an ill-posed inference problem due to the absence of direct measurements for the transverse velocity component *v*_*θ*_, rendering the observation matrix *A* in equation (1) rank-deficient. This challenge is analogous to optical flow estimation in directions perpendicular to image gradients [32]. The divergence operator—VFM’s principal physical constraint—is itself singular and insufficient for regularization unless supplemented with proper smoothing and boundary conditions. This section analyzes the the B-VFM precision matrix, which is the equivalent of the normal matrix *Q*^*⊤*^*Q* in least-squares formulations of the form 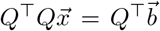 [4]. We examine the structure of its inverse to formalize the ill-posedness of VFM, and assess its impact on inference and uncertainty. We then derive the null space of the mass conservation constraint to characterize VFM artifacts and analyze if and how boundary conditions and smoothing suppress these artifacts.

#### 3.1.1 Structure of the VFM Covariance Matrix

Here, we investigate the structure of the log-conditional posterior Hessian inverse for VFM, which is proportional to the covariance matrix at the MAP estimate, 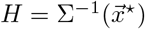. To simplify our analysis, we start with the block decomposition of *H*

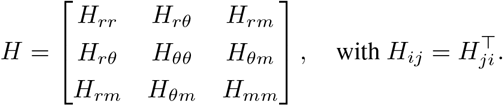

See §SI2 in the Appendix for an explicit derivation of this Hessian matrix. To keep the analysis concise, we further assume we are working with a good-quality image with low Doppler and mask observation errors, i.e., Σ_*V*_, Σ_*M*_ *≪* 1, so that

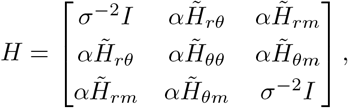

where *σ*^2^ *∼* Σ_*V*_ *∼* Σ_*M*_, *α* represents a generic strength of the VFM priors, i.e., the hyperparameters, and the tilde is used to indicate that the Hessian blocks have been normalized to have a similar norm as the identity matrix *I*. Using a first-order Neumann expansion, the inverse of *H* is approximated as (§SI2 in the Appendix)

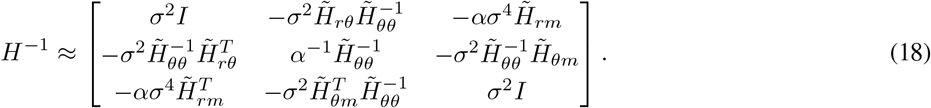

In contrast, a data assimilation problem with observation in both *v*_*r*_ and *v*_*θ*_ would have a precision matrix of the form

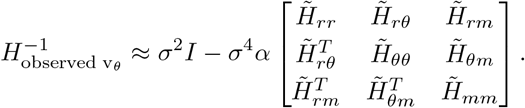

This difference has an important implication. The Neumann approximation for *H*^*−*1^ breaks down and *H* becomes singular when *α ≪* 1, a reminder of the impossibility of inferring *v*_*θ*_ without imposing physical constraints. This issue does not arise in 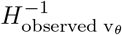

We also note that only the (*r, m*) blocks in *H*^*−*1^ are quadratic in *σ*, indicating that Doppler and mask are more weakly coupled than other variables. The coupling block 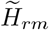 arises from the interdependence between mask and Doppler velocity in the boundary conditions (equation 8). Detailed analysis indicates that this coupling is activated at pixels where the mask gradient and radial velocity fail to satisfy the free-slip boundary condition prior, and that its magnitude scales as *O*(Σ_*V*_ Σ_*M*_ *α*_*BC*_), reflecting weaker coupling when either the Doppler velocity or LV mask are well observed (i.e., low error), and stronger coupling when the boundary condition prior is more strongly enforced. In traditional VFM methods, there is no possible coupling since the mask is not a latent variable, which effectively sets Σ_*M*_ = 0. Therefore, these methods cannot correct segmentation errors using velocity data.

The diagonal of *H*^*−*1^ provides an estimate of the posterior uncertainty in the latent variables assuming low hyperparameter variance. In the limit of small measurement noise (*Σ* ≪ 1), the first-order Neumann series indicates that the uncertainty in *v*_*r*_ and *m* is mostly dictated by their measurement noise. In contrast, the uncertainty in *v*_*θ*_ depends entirely on the hyperparameters and the inverse of the central block, 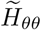, which encodes the *θ*-component of the mass conservation, boundary conditions, and smoothing priors.

#### 3.1.2 Effect of Blood-Domain Geometry and Priors on VFM Ill-Posedness

Equation (18) shows that, at leading order, the regularity of *H* depends critically on the regularity of the central block 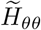. Here, we analyze the regularity of this matrix by finding the singular eigenvectors of the *θ*-component of the discretized divergence operator (i.e., spurious modes that could be summed to *v*_*θ*_ without breaking mass conservation) and computing how the boundary conditions and smoothing priors modify the singular eigenvalue (i.e., how those priors damp the strength of the spurious modes).

To simplify our derivations, we will work first in 1D and will later extend key results to the 2D case relevant to VFM. Let 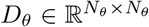 be the discretized derivative operator ∂*/*∂*θ* with periodic boundary conditions. This operator’s nullspace is defined by the *N*_*θ*_ *×* 1 vector 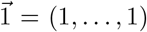, yielding 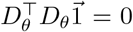. This vector also belongs to the nullspace of any 1D smoothing operator, so it can only be supressed by boundary conditions. Enforcing a no-flux boundary condition 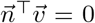 together with mass conservation leads to

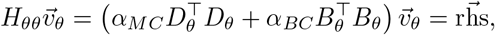

where 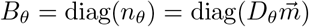 is a diagonal matrix encoding the *θ*-component of the normal vector to the mask, while *α*_*MC*_ and *α*_*BC*_ are the hyperparameters controlling each penalty’s strength. Thus, imposing boundary conditions can be seen as a rank update to the precision matrix. Its effect on the null eigenvalue of *D*_*θ*_ corresponding to the constant vector 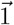 is determined by computing

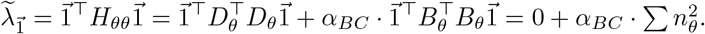

Extending this analysis to 2D is straightforward because the *N*_*r*_ null eigenvectors of *D*_*θ*_ are orthogonal to each other. Therefore, we can study each eigenvector separately, similar to the 1D case. These eigenvectors are denoted 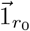, and have a constant value in *θ* along the radial arc *r* = *r*_0_, and vanish for *r ≠ r*_0_. However, their eigenvalues get an additional update from the smoothing prior because the eigenvectors 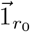_*r*_ do not belong to the nullspace of the discretized Laplacian operator, i.e.,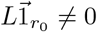. Assuming second-order centered finite differences, the regularized 2D eigenvalues become:

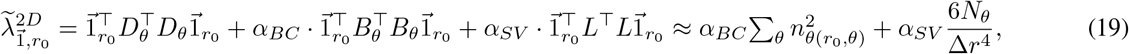

where *α*_*SV*_ is the smoothing penalty hyperparameter, Δ*r* is the polar mesh spacing in the radial direction and we have made the approximation *r*_0_ *≫* Δ*r*.

To complete the 2D analysis, we consider the intersection of the nullspaces of the mass conservation and smoothing operators, where regularization can only come from boundary conditions. This space has dimension one and is spanned by a *N*_*r*_ *× N*_*θ*_ vector full of ones, representing a constant function in both the *r* and *θ* directions, 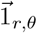. For this vector, it is straightforward to obtain that 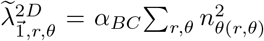, guaranteed to be positive as long as there is one pixel in the entire domain with *n*_*θ*_ *≠* 0. Therefore, this degeneracy should not be concerning in VFM.

Finally, it is worth studying how the singular eigenvalues of mass conservation would be modified when no-slip instead of free-slip boundary conditions were enforced. In that case, *n*_*θ*_ should be replaced by an indicator function *δ*_*LV*_ (*r, θ*) that labels endocardial pixels where flow velocity matches endocardial velocity. The modified eigenvalues would then be

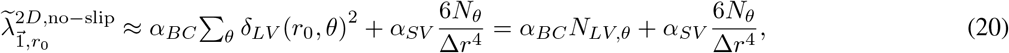

where *N*_*LV,θ*_ is the number of pixels segmented as endocardial at *r* = *r*_0_. Thus, the regularity of the cross-beam velocity inference is governed by the geometry of the boundary constraints and, when needed, by smoothing imposed during inference.

In summary, when smoothing is omitted during inference or is small (i.e., *α*_*SV*_ *≪* 1), the well posedness of VFM cannot be guaranteed and can be predicted from the geometry of the segmented mask (Figure **??**). Under free-slip conditions, VFM is well posed only if, at each constant-*r* arc, the mask boundary is not completely aligned with the *θ*-direction. In that case, the transverse normal component *n*_*θ*_ is zero and the term multiplying *α*_*BC*_ in equation (19) vanishes. For no-slip conditions, regularity requires that there are no holes through the entire LV mask at any constant-*r* arc, so that the number of segmented pixels *N*_*LV,θ*_ is nonzero and the term multiplying *α*_*BC*_ in equation (20) remains active. When the geometric criteria are not satisfied, *v*_*θ*_ becomes undetermined by constant along any affected arc, producing band-like artifacts in the reconstructed velocity field. Such artifacts can be observed in VFM flow maps in the literature [25, 35].

### 3.2 B-VFM Performance for 2D Vortex Flow

This section evaluates the performance of B-VFM and compares it with traditional approaches using as a benchmark the flow generated by the Lamb–Chaplygin (LC) dipole introduced in §2.3. The LC flow field is decomposed into radial and transverse components; the transverse component is discarded, and the radial component is taken as the observed Doppler field *V*. The blood domain *M* is defined by the circular dividing streamline that encloses the recirculating fluid within the LC vortices. We place particular emphasis on examining the behavior of B-VFM under non-ideal conditions, where the input data 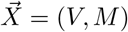 may include spurious Doppler reflections or inaccuracies in blood-domain segmentation. In these scenarios, the ideal noiseless LC dipole serves as our benchmark flow, while the least-squares formulation iVFM [4] provides the reference VFM method for comparison. When comparing with this method, we will use the nomenclature iVFM[*λ*_*MC*_,*λ*_*BC*_,*λ*_*SM*_], where *λ*_*MC*_, *λ*_*BC*_, and *λ*_*SM*_ are the weights of the mass-conservation, boundary conditions, and smoothing penalties relative to the Doppler misfit term.

#### 3.2.1 Ideal Benchmark: Noiseless 2D Flow with Perfect Segmentation

We first consider the ideal benchmark, in which the flow field is noiseless, strictly planar, and perfectly segmented. Under these conditions, both B-VFM and iVFM are expected to recover the flow field up to numerical discretization error. Because the input data contained no uncertainty, i.e.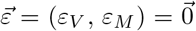. the corresponding precision parameters 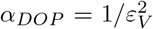 and 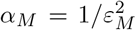 would be formally infinite. In practice, we retained these as hyperparameters to be inferred. This approach allowed the sampler to determine the relative weighting of the Doppler and segmentation misfit terms and provided an automatic scaling among the different model residuals.

We ran ten MCMC chains concurrently for 200 iterations for each case reported in this manuscript. We used MATLAB R2022b on a Desktop computer running Linux version 6.8.0-138-generic equipped with 28 CPU Intel(R) Xeon(R) w7-3465X cores, with 76MB Cache Memory per core, and 128 total GB RAM. Each run took approximately 30 minutes. Inspection of the hyperparameter trajectories for ten different chains (Figure 4*A*) shows robust convergence regardless of the range of initial conditions. Based on these trajectories, we set the burn-in period to be *it*^*b*^ = 80 iterations in this case. After burn-in, the marginal posterior distributions of the hyperparameters, 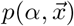 closely overlapped across chains (Figure 4*B*), providing strong evidence that the chains were sampling the same stationary posterior distribution. The posterior correlations revealed coupling among several model terms. In particular, *α*_DOP_ was positively correlated with both *α*_Seg_ and *α*_MC_, whereas *α*_BC_ was negatively correlated with *α*_DOP_, *α*_Seg_, and *α*_MC_. By contrast, the smoothing hyperparameters were largely independent of the remaining hyperparameters. These results suggest that the physical constraints cross-talked with both the input mask and Doppler data during Gibbs sampling.

**Figure 4:**
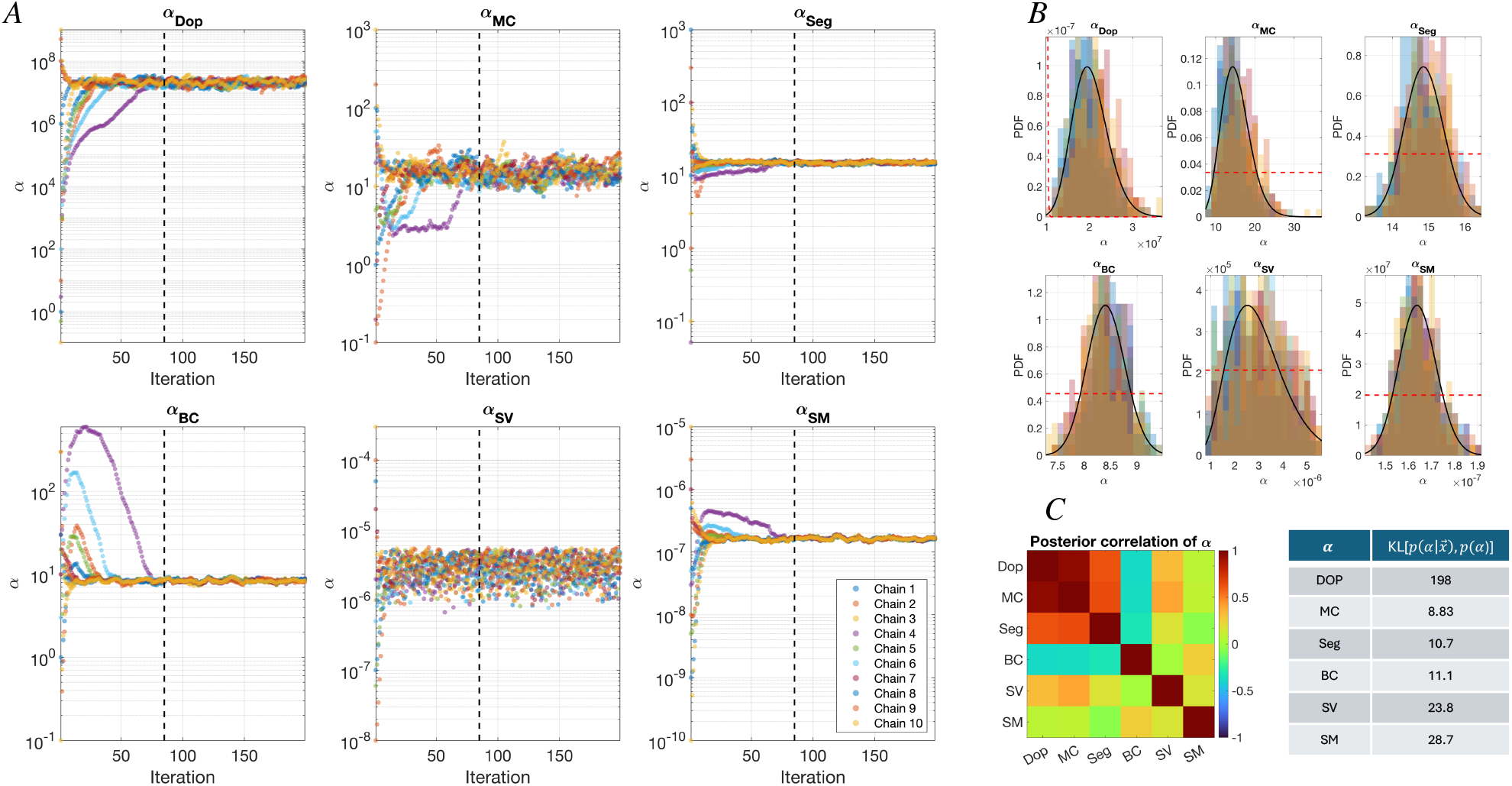
B-VFM Hierarchical Bayesian Sampler for the ideal LC dipole case. *A*) Ten hyperparameter Markov chains; the dashed vertical line separates the burn and collection pieces of the chains. *B*) Overlaid posterior probability distributions of the hyperparameters, 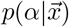, for the ten chains.——, Gamma distribution fit to 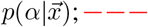, −− −, non-informative hyperparameter prior *p*(*α*) with shape and rate parameters 1 and 10^*−*5^. *C*) Posterior corelation between hyperparameters and Kullback-Leibler divergence of the hyperparameter posterior and prior distributions.

To estimate how much information was gained from the data, we computed the Kullback-Liebler divergence between the prior and marginal posterior distributions of hyperparameters, 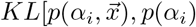 (Table in Figure 4*C*). This analysis revealed that the Doppler weight underwent the largest posterior update relative to its prior, followed by the mask- and velocity-smoothing weights. The boundary-condition, segmentation, and mass-conservation weights exhibited smaller, although still appreciable, updates.

We next examined the posterior mean the velocity and mask fields obtained from the Gibbs samples retained after burn-in (Figure 5*A*). As expected, B-VFM reconstructed *v*_*θ*_ in excellent agreement with both the LC-dipole ground truth and the iVFM[1, 0.1, 10^*−*8^] solution. B-VFM also accurately recovered the mask *m* while providing spatially resolved estimates of the posterior variances of *v*_*r*_, *v*_*θ*_, and *m*.

**Figure 5:**
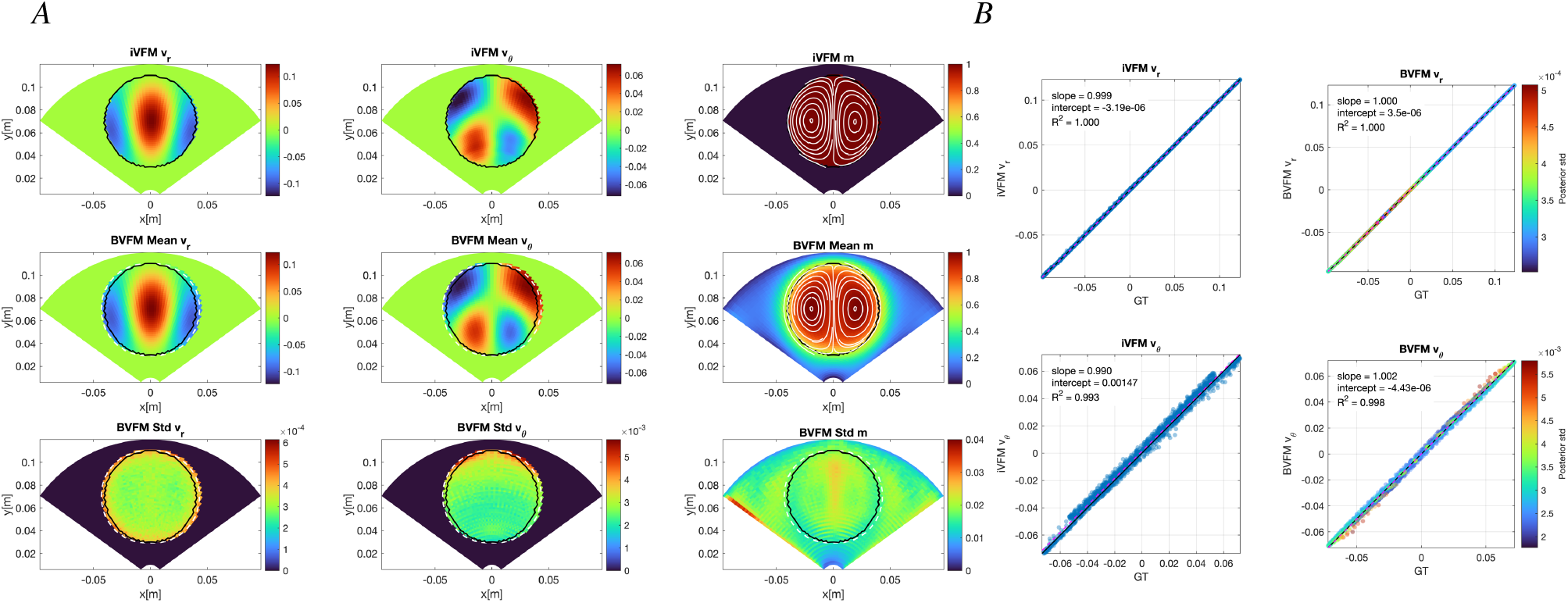
Reconstruction of the ideal, noiseless Lamb–Chaplygin (LC) dipole. *A*) Spatial distributions of *v*_*r*_, *v*_*θ*_, and the mask *m* with superimposed flow streamlines (left to right). Top row: iVFM [1, 0.1, 10^*−*8^] reconstruction and reference binary mask; middle row: B-VFM posterior means; bottom row: B-VFM posterior standard deviations. The solid black and white dashed contours denote respectively the boundaries of the reference and inferred mask, defined by *M* = 0.5 and *m* = 0.5 respectively. *B*) Pointwise comparisons of the ground-truth (GT) and reconstructed velocity components (top: *v*_*r*_, bottom: *v*_*θ*_; left: iVFM[1,0.1,10^*−*8^], right: B-VFM). B-VFM points are colored according to their posterior standard deviation. ——,identity line; 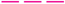, least-squares regression line. Velocity and their posterior standard deviations are given in m,s^*−*1^, and the mask is dimensionless.

The inferred mask achieved a Dice similarity coefficient of 0.94 relative to the reference mask. Its only appreciable discrepancy was a slight widening near the equator of the vortex, as indicated by the separation between the black and white dashed contours in the the second and third rows of Figure 5*A*. The inferred mask exhibited a weak tendency to develop a lobed structure within the vortex. This behavior can be attributed to the boundary-condition prior, 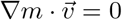, which favors masks that are constant along the flow streamlines (white lines in the mask panels of Figure 5*A*). This minor artifact did not measurably affect the velocity reconstruction. Setting a hard constraint at points well inside or well outside the vortex (see §2.2.3) eliminated the lobed structure and improved the Dice coefficient to 0.98, confining the posterior variance of *m* to the vicinity of the vortex boundary, while leaving the remaining results essentially unchanged (see Figure SI1 in the Appendix).

The posterior standard deviations were uniformly small, as expected for noiseless input data and a flow for which planar mass conservation holds exactly. The bottom row of Figure 5*A* shows that 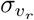 was approximately one order of magnitude smaller than 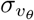, consistent with the analysis presented in the previous section. Also worth noting, *σ*_*m*_ developed a modest ridge along the axis separating the two lobes described above.

Figure 5B presents pointwise comparisons between the ground-truth and inferred values of *v*_*r*_ and *v*_*θ*_, confirming the excellent performance of both B-VFM and iVFM for the LC-dipole flow. For B-VFM, the coefficients of determination were *R*^2^ = 1.000 for *v*_*r*_ and *R*^2^ = 0.998 for *v*_*θ*_. The corresponding values for iVFM [1, 0.1, 10^*−*8^] were *R*^2^ = 1.000 and *R*^2^ = 0.993, respectively. The slopes of the linear fits for *v*_*r*_ were 1.000 for both methods. For *v*_*θ*_, the slopes were 1.001 with B-VFM and 0.990 with iVFM. The fitted intercepts were negligible in all cases. Increasing the smoothing level in iVFM produced more regular *v*_*θ*_ patterns but slightly decreased the pointwise accuracy (see iVFM [1, 0.1, 10^*−*6^] results in Figure SI1 in the Appendix).

#### 3.2.2 Color-Doppler Artifact Removal

When implanted metallic objects (mechanical prosthetic valves, implanted device leads, etc) interact with ultrasound, they can generate twinkling artifacts, a rapidly fluctuating, random mixture of red and blue on color Doppler within the angular sector in the acoustic shadow of the object. These artifacts may mimic blood flow where none exists or obscure the true velocity signal. To evaluate B-VFM under similar conditions, we selected a fan-shaped region within the Doppler sector that overlapped the flow domain. Within this region, we replaced the Doppler signal with random values uniformly distributed between *±U*_*max*_, where *U*_*max*_ = *max*(|*V* |). Additionally, we substituted the spatially uniform data reliability, Σ_*V*_ = *α*_*DOP*_ *I*, by a diagonal matrix diag(*σ*_*V*_) where the *σ*_*V*_ field reflects the local signal-to-noise ratio. Specifically, we set *σ*_*V*_ within the artifact region to be 10-fold higher than in the rest of the domain, informing the model to place less confidence in the corrupted Doppler data.

Inference was performed using the Gibbs sampler (Algorithm 1) with ten independent chains as in the ideal benchmark. All chains converged to similar posterior estimates, demonstrating robust and reproducible inference (Figure 6*A*). The inferred hyperparameters were largely consistent with those of the ideal benchmark (Figure 4), except for *α*_*DOP*_, which was markedly lower in the presence of the artifact. Accordingly, the KL divergence between the posterior and hyperprior distributions of *α*_*DOP*_ also decreased, indicating that B-VFM adapted its model for color Doppler to be less informative than in the ideal case, consistent with the presence of corrupted Doppler data.

**Figure 6:**
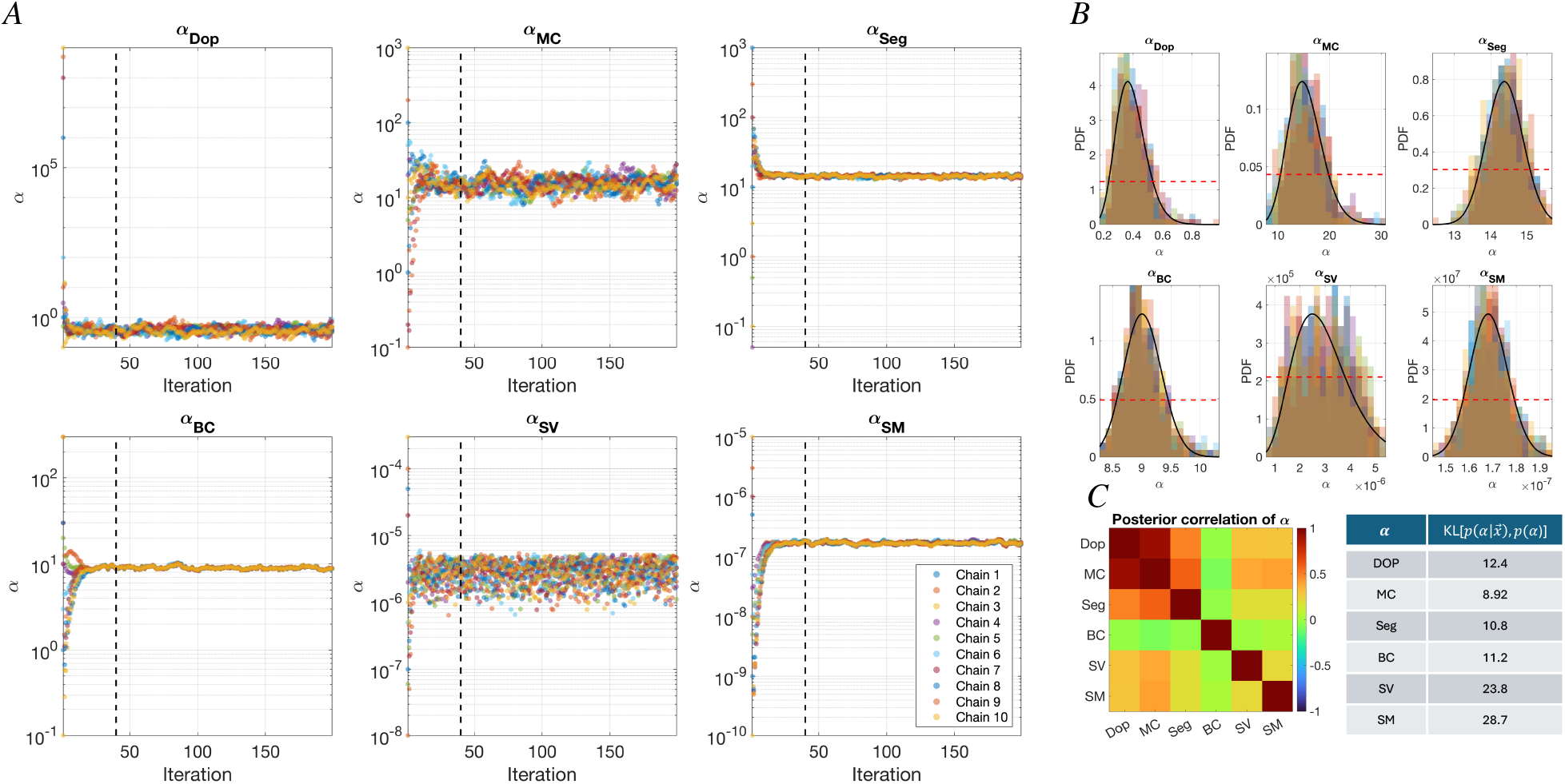
B-VFM Hierarchical Bayesian Sampler for an LC dipole with a color Doppler (see fig. 7*A*) Ten hyperparameter Markov chains; the dashed vertical line separates the burn and collection pieces of the chains. *B*) Overlaid posterior probability distributions of the hyperparameters, 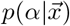, for the ten chains. ——, Gamma distribution fit to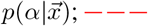, non-informative hyperparameter prior *p*(*α*) with shape and rate parameters 1 and 10^*−*5^. *C*) Posterior corelation between hyperparameters and Kullback-Leibler divergence of the hyperparameter posterior and prior distributions.

Figure 7*A* compares the expected velocity and mask fields inferred by B-VFM with those obtained using iVFM[1,0.1,10^*−*8^].

**Figure 7:**
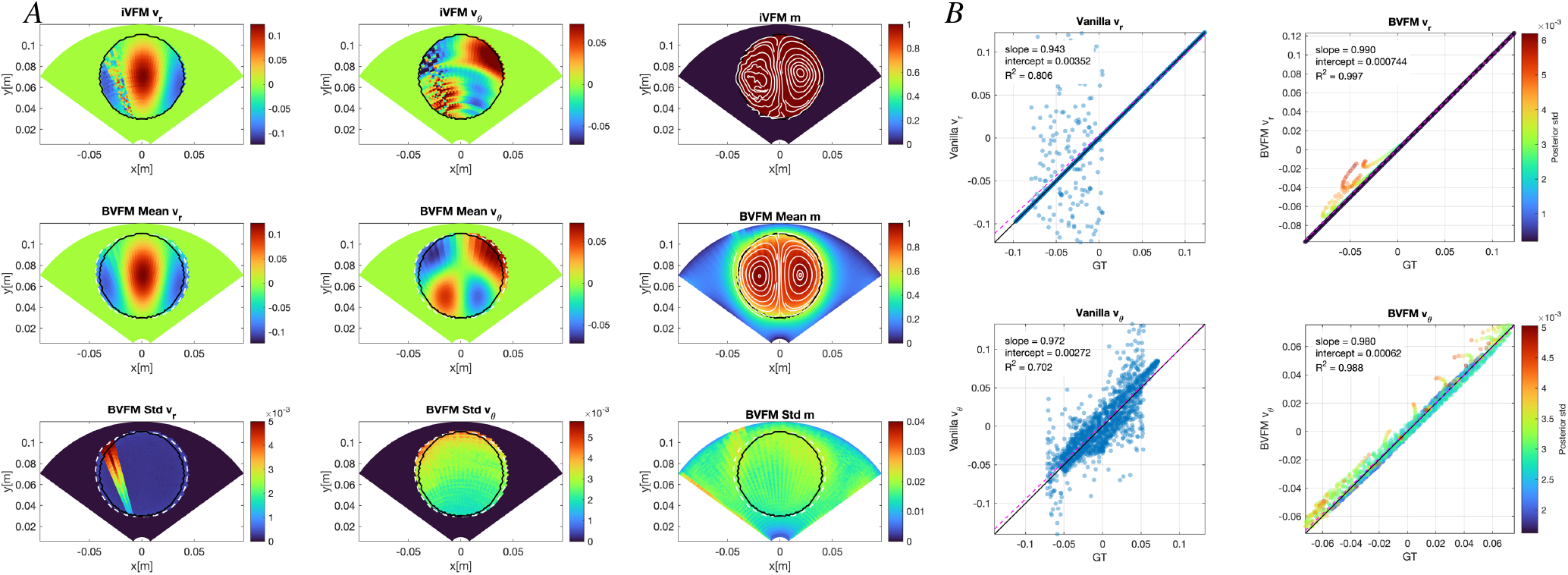
Reconstruction of a Lamb–Chaplygin (LC) dipole with a twinkling Doppler artifact. *A*) Spatial distributions of *v*_*r*_, *v*_*θ*_, and the mask *m* with superimposed flow streamlines (left to right). Top row: iVFM [1, 0.1, 10^*−*8^] reconstruction and reference binary mask; middle row: B-VFM posterior means; bottom row: B-VFM posterior standard deviations. The solid black and white dashed contours denote respectively the boundaries of the reference and inferred mask, defined by *M* = 0.5 and *m* = 0.5 respectively. *B*) Pointwise comparisons of the ground-truth (GT) and reconstructed velocity components (top: *v*_*r*_, bottom: *v*_*θ*_; left: iVFM[1,0.1,10^*−*8^], right: B-VFM). B-VFM points are colored according to their posterior standard deviation.——, identity line; 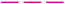, least-squares regression line. Velocity and their posterior standard deviations are given in m,s^*−*1^, and the mask is dimensionless.

B-VFM reconstructed smooth, continuous velocity patterns across the twinkling-artifact region while largely preserving the symmetric dipole structure and circular segmentation mask. In contrast, iVFM[1,0.1,10^*−*8^] reproduced the Doppler fluctuations within the fan, generating similar disturbances in *v*_*θ*_, and exciting the banded instability associated with the 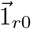 null vector of *D*_*θ*_ described in §3.1.2. These banded instabilities poluted *v*_*θ*_ in the entire flow domain. These artifacts can be effectively removed by increasing the smoothing parameter in iVFM at the cost of decreasing the pointwise accuracy of *v*_*θ*_ (see iVFM[1,0.1,10^*−*4^] results in Figure SI2 of the Appendix).

By accounting for spatial variations in input uncertainty, B-VFM placed less weight on *V* and greater weight on the priors where Σ_*V*_ was elevated. The posterior variance 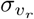 was correspondingly higher within the artifact fan (Figure 7*A*, bottom row). This uncertainty propagated to *v*_*θ*_ and *m* through the Hessian cross-terms and posterior hyperparameter distributions, especially at higher depths.

Scatter plots comparing the inferred and ground-truth velocities (Figure 7*B*) confirmed the superior accuracy of B-VFM. Pointwise comparisons yielded coefficients of determination of *R*^2^ = 0.997 for *v*_*r*_ and *R*^2^ = 0.988 for *v*_*θ*_. The corresponding linear-fit slopes were 0.990 and 0.979, with negligible intercepts. The inferred mask achieved a Dice score of 0.94, similiar to the ideal case. Moreover, coloring the data points by posterior uncertainty showed that B-VFM reliably identified the least reliable velocity estimates. In comparison, iVFM[1,0.1,10^*−*8^] yielded *R*^2^ = 0.806 for *v*_*r*_ and *R*^2^ = 0.702 for *v*_*θ*_, with linear-fit slopes of 0.943 and 0.972, respectively. The *v*_*θ*_ scatter plot further showed that errors originating within the artifact fan propagated throughout the field, whereas errors in the Doppler component, *v*_*r*_, remained largely confined to the fan, consistent with the reconstructed fields in Figure 7*A*.

Applying hard mask constraints produced results similar to those of unconstrained B-VFM (Figure SI2 in the Appendix), while modestly improving segmentation accuracy (Dice score, 0.98) and concentrating the posterior uncertainty in *v*_*θ*_ and *m* more closely around the Doppler-artifact region.

#### 3.2.3 Segmentation Correction

Clinical segmentation of the LV wall is often imperfect, and such errors can substantially degrade VFM reconstructions by introducing inconsistencies between the color-Doppler data (*V*) and the prescribed boundary conditions. Traditional VFM methods treat the segmentation as deterministic, attributing such inconsistencies to errors in *V* and thereby biasing the inference of *v*_*θ*_ [22, 33, 49, 52, 76]. Least-squares formulations with tunable weights [4] can partially mitigate this mismatch, but with a prescribed segmentation, an inaccurate boundary creates an unavoidable tradeoff: decreasing the weight of boundary-condition penalties weakens the physical fidelity of the inferred flow field, and increasing it propagates the segmentation error into the inferred velocity. Tuning penalty weights cannot correct the mask itself, motivating the joint inference of velocity and segmentation in B-VFM.

B-VFM addresses this limitation by treating the segmentation mask probabilistically, allowing inconsistencies with color-Doppler data to be resolved through the priors. To illustrate this capability, we created a dent in the observed *M* and set σ_*M*_ = *α*_*Seg*_ diag(*σ*_*M*_), with *σ*_*M*_ = 1 throughout the domain except in the dented region, where we set *σ*_*M*_ = 10 to reflect lower confidence on *M* within that region.

With these input data, the Gibbs sampler behaved similarly to the ideal case in terms of rapid convergence. Interestingly, posterior hyperparameter values indicated that mass conservation and velocity regularization received greater weight, while boundary conditions were downweighted slightly in comparison to the ideal case (Figure 8).

**Figure 8:**
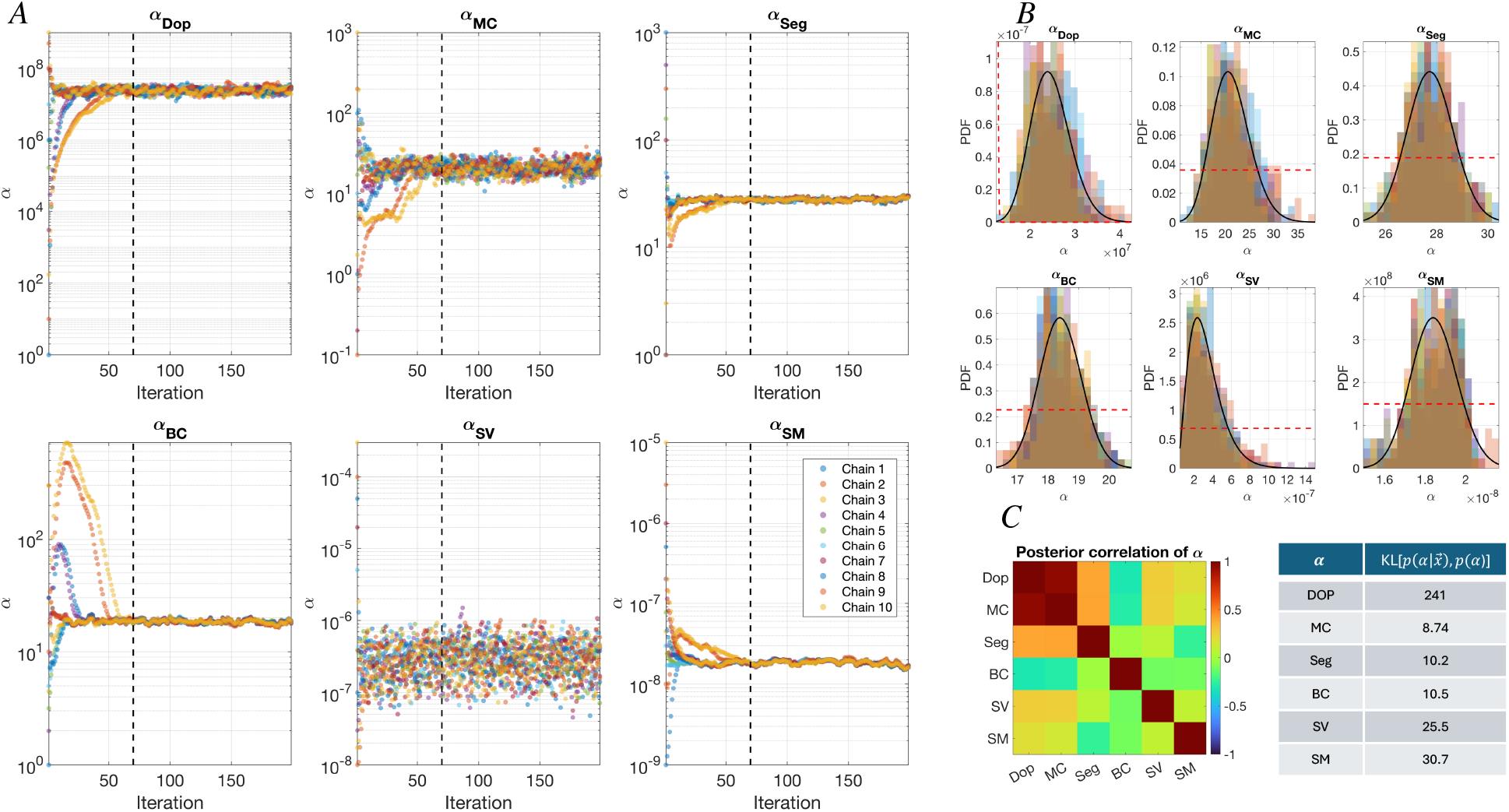
B-VFM Hierarchical Bayesian Sampler for an LC dipole with a mask defect (see fig 9*A*) Ten hyperparameter Markov chains; the dashed vertical line separates the burn and collection pieces of the chains. *B*) Overlaid posterior probability distributions of the hyperparameters, 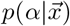, for the ten chains.— —, Gamma distribution fit to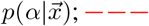;−−−, non-informative hyperparameter prior *p*(*α*) with shape and rate parameters 1 and 10^*−*5^. *C*) Posterior corelation between hyperparameters and Kullback-Leibler divergence of the hyperparameter posterior and prior distributions.

Figure 9*A* shows that B-VFM recovered velocity fields in close agreement with the ground truth and inferred a mask *m* in which the artificial dent present in the input mask *M* was largely corrected (dice score of 0.98 with the reference circular mask). Thus, B-VFM used the modeled uncertainty in the wall mask to reconcile local inconsistencies between the observations and the physical priors. In contrast, iVFM[1, 0.1, 10^*−*8^] treated the erroneous mask as exact, imposing spurious boundary conditions that distorted *v*_*θ*_ along radial arcs intersecting the dent. This perturbation worsens as the weight of the boundary condition constraint is increased, requiring significant smoothing to prevent banded artifacts, and overall decreasing inference accuracy (see Figure SI3 in the Appendix).

**Figure 9:**
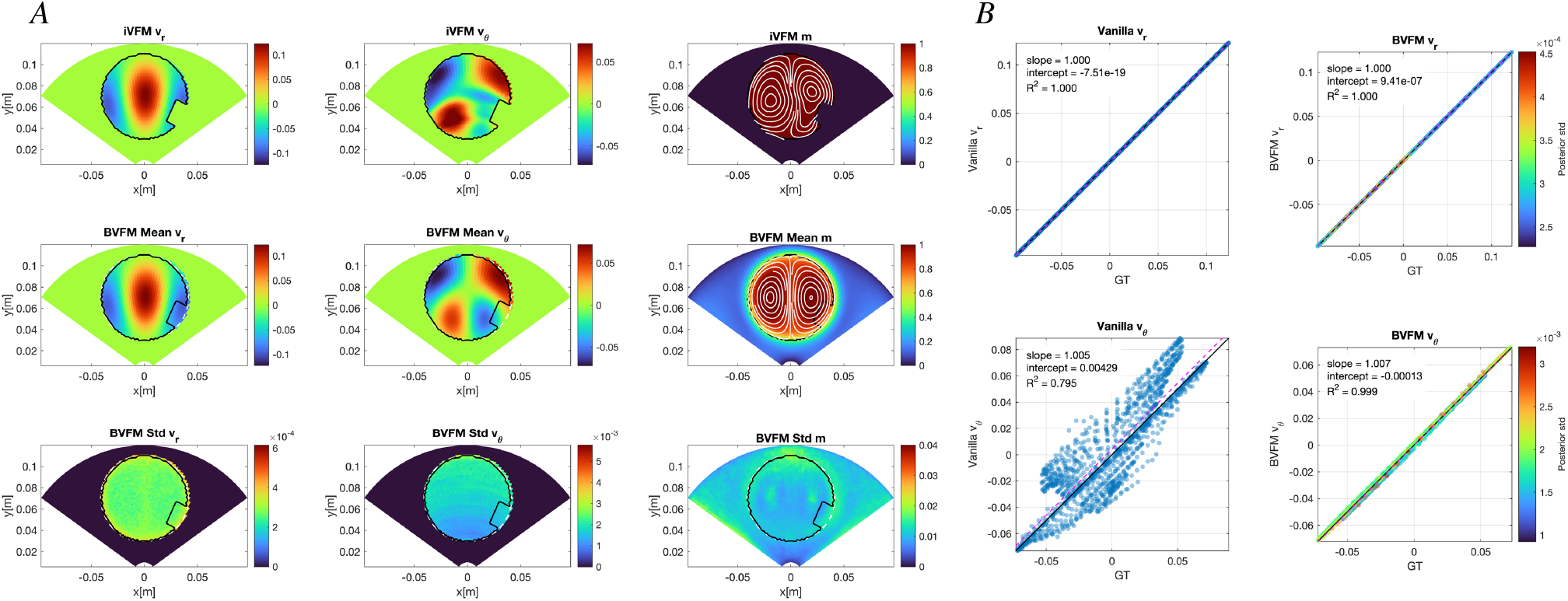
Reconstruction of a Lamb–Chaplygin (LC) dipole with a twinkling Doppler artifact. *A*) Spatial distributions of *v*_*r*_, *v*_*θ*_, and the mask *m* with superimposed flow streamlines (left to right). Top row: iVFM [1, 0.1, 10^*−*8^] reconstruction and reference binary mask; middle row: B-VFM posterior means; bottom row: B-VFM posterior standard deviations. The solid black and white dashed contours denote respectively the boundaries of the reference and inferred mask, defined by *M* = 0.5 and *m* = 0.5 respectively. *B*) Pointwise comparisons of the ground-truth (GT) and reconstructed velocity components (top: *v*_*r*_, bottom: *v*_*θ*_; left: iVFM[1,0.1,10^*−*8^], right: B-VFM). B-VFM points are colored according to their posterior standard deviation.——, identity line; 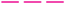, least-squares regression line. Velocity and their posterior standard deviations are given in m,s^*−*1^, and the mask is dimensionless.

The posterior uncertainty maps (Figure 9*A*, bottom row) closely resembled those obtained for the ideal benchmark. Notably, the posterior standard deviation of *m* was not elevated near the dent. Thus, although the input mask *M* was assigned substantial uncertainty in this region, B-VFM corrected the defect with high posterior precision, indicating that the Doppler observations and physical priors strongly constrained the local mask geometry.

To examine this interpretation, we repeated the analysis using the same dented input mask but without locally increasing Σ_*M*_, thereby treating the erroneous mask values within the dent as being as precise as those elsewhere. To facilitate convergence, we also imposed hard constraints on *m* at points well inside and outside the mask. Results are shown in Figure SI3 of the Appendix. B-VFM again corrected most of the dent, although a mild erosion remained near the defect and a slight dilation appeared on the opposite side of the domain at comparable radii, yielding a Dice score of 0.96. More importantly, the conflict between the now high-confidence *M* and the physical priors produced a marked increase in the posterior standard deviation of *m* within the dented region. This uncertainty also propagated into *v*_*θ*_ along the characteristic radial-band pattern.

Pointwise comparisons showed excellent agreement between the B-VFM-inferred and ground-truth velocities: both *v*_*r*_ and *v*_*θ*_ yielded *R*^2^ = 1.000 and linear-regression slopes of 1.000 (Figure 9*B*). iVFM[1, 0.1, 10^*−*8^] also produced slopes close to unity (1.000 and 1.005 for *v*_*r*_ and *v*_*θ*_, respectively), but its transverse-velocity estimates exhibited substantially greater scatter (*R*^2^ = 0.795).

### 3.3 B-VFM Performance for 3D Synthetic Vortex Flow

Having established B-VFM performance for the planar LC dipole, we next challenged its planar-flow assumption using the family of Hicks–Moffatt (HM) spherical vortices described in §2.3. These vortices are incompressible in 3D but their meridional-plane velocity fields are not divergence-free on that plane. They therefore provide a controlled test of how B-VFM balances an inconsistent mass-conservation prior against the remaining priors and the observed Doppler data.

We first considered Hill’s spherical vortex, the member of the HM family corresponding to the dimensionless wavenumber *κ* = 0. Without hard mask constraints, B-VFM was sensitive to hyperparameter initialization and, for some initial values, failed to reach a physically plausible solution. Hard mask constraints substantially improved convergence robustness: all constrained chains converged, and whenever the unconstrained formulation converged successfully, the constrained and unconstrained formulations produced similar solutions. Accordingly, Figure 10 presents results from ten constrained chains whose hyperparameters were initialized randomly over broad prescribed ranges.

**Figure 10:**
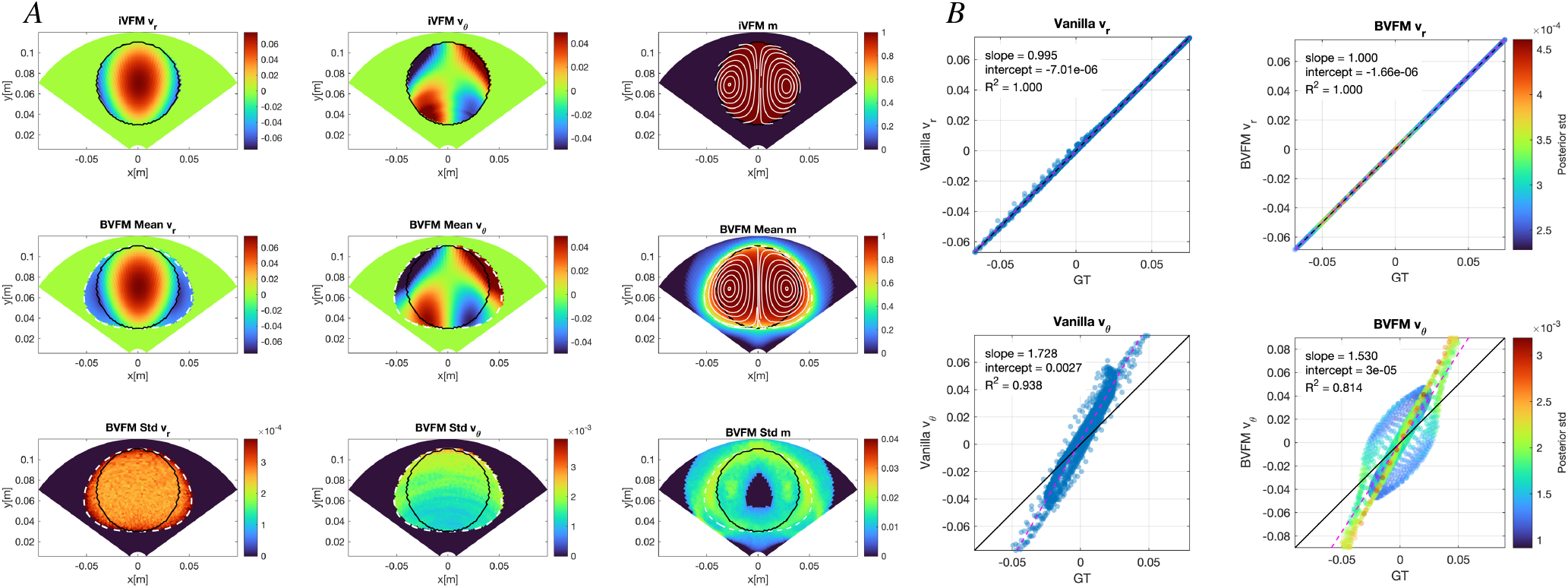
Reconstruction of a Hill’s spherical vortex flow. *A*) Spatial distributions of *v*_*r*_, *v*_*θ*_, and the mask *m* with superimposed flow streamlines (left to right). Top row: iVFM [1, 10^4^, 10^*−*6^] reconstruction and reference binary mask; middle row: B-VFM posterior means; bottom row: B-VFM posterior standard deviations. The solid black and white dashed contours denote respectively the boundaries of the reference and inferred mask, defined by *M* = 0.5 and *m* = 0.5 respectively. *B*) Pointwise comparisons of the ground-truth (GT) and reconstructed velocity components (top: *v*_*r*_, bottom: *v*_*θ*_; left: iVFM[1,0.1,10^*−*8^], right: B-VFM). B-VFM points are colored according to their posterior standard deviation.——, identity line; 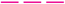, least-squares regression line. Velocity and their posterior standard deviations are given in m,s^*−*1^, and the mask is dimensionless.

B-VFM expanded the inferred mask beyond the reference boundary, yielding a Dice score of 0.86. This expansion allowed the reconstruction to accommodate the observed flow while satisfying the no-penetration boundary condition, as indicated by streamlines that remained approximately tangent to the inferred mask boundary. While iVFM[1, 0.1, 10^*−*8^] retained the prescribed mask, it produced streamlines that crossed its boundary (compare the first and second rows of Figure 10*A*). Accordingly, the boundary-condition residual was substantially larger for iVFM (Figure SI4 in the Appendix). Increasing the boundary-condition weight *λ*_BC_ in iVFM improved the alignment between the streamlines and the prescribed mask boundary. In our tests, however, this required stronger velocity smoothing, which excessively attenuated the reconstructed Doppler and the inferred *v*_*θ*_ fields (not shown). Despite these differences, B-VFM and iVFM recovered *v*_*θ*_ fields with similar overall spatial structure. Their planar mass-conservation residual, 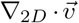, also exhibited similar spatial patterns, although its magnitude was significantly larger for B-VFM (Figure 10 in the Appendix).

Pointwise comparisons showed that both methods accurately reproduced the ground-truth radial velocity (Figure 10*B*). For B-VFM and iVFM, respectively, the comparisons yielded *R*^2^ = 1.000 and 0.984, with linear-fit slopes of 1.000 and 0.995. Enforcing planar mass-conservation to the meridional section of the 3D Hill flow caused both methods to overestimate *v*_*θ*_, yielding slopes of 1.530 for B-VFM and 1.728 for iVFM. Although B-VFM reduced this bias, it exhibited greater pointwise scatter than iVFM (*R*^2^ = 0.814 and 0.938, respectively). Overall, both B-VFM and iVFM showed reduced transverse-velocity accuracy for the three-dimensional Hill flow, underscoring the intrinsic limitations of planar VFM when out-of-plane transport violates 2D mass conservation.

The posterior uncertainty maps from B-VFM localized the principal discrepancies from the ground truth. The largest values of 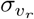 and *σ*_*m*_ occurred in regions where the inferred mask extended beyond the reference mask, whereas the spatial distribution of 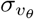 resembled that of the planar mass-conservation residual (Figure 10 in the Appendix).

#### 3.3.1 Effect of the ensemble-informed mass-conservation prior

We next investigated whether BVFM accuracy could improve by informing the 2D mass-conservation prior with statistics obtained from a family of related 3D flows. Assuming that empirical information about the mean and covariance of the in-plane mass-conservation residual is available, one could formulate a data-driven BVFM by recasting the the mass conservation prior (equation 3) as

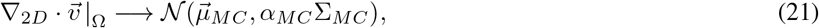

where 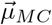 replaces zero and a full covariance matrix Σ_*MC*_ replaces the identity matrix. In clinical applications, these statistics could be learned from 2D velocity fields acquired using phase-contrast MRI or a different method. Here, we estimated them from an ensemble of 40 HM vortices with uniformly spaced values of *κ* between 0 and 5.5, covering a range of different vortex dipole shapes and intensities, as described in § 2.3.1 (Figure 2).

Figure 11*A* presents results from ten independent B-VFM chains using the fully data-driven mass-conservation prior, incorporating both the learned mean 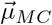and covariance Σ_*MC*_. The reconstructed flow pattern closely matched the ground truth, while the inferred mask showed minimal distortion and achieved a Dice score of 0.98. Moreover, the streamlines remained approximately tangent to the inferred mask boundary, indicating that the boundary conditions were well satisfied. Scatter plots (Figure 11*B*) show that the Doppler velocity *v*_*r*_ was recovered exactly (linear-fit slope 1.000, *R*^2^ = 1.000), while *v*_*θ*_ was reconstructed with a slope of 0.935 and *R*^2^ = 0.882. The characteristic nonzero planar divergence was also accurately reproduced, with a slope of 0.820 and *R*^2^ = 0.806. Thus, combining the learned mean and covariance enabled B-VFM to reproduce both the velocity field and its 3D contribution to the planar mass-conservation residual, substantially mitigating the limitations of the planar-flow assumption.

**Figure 11:**
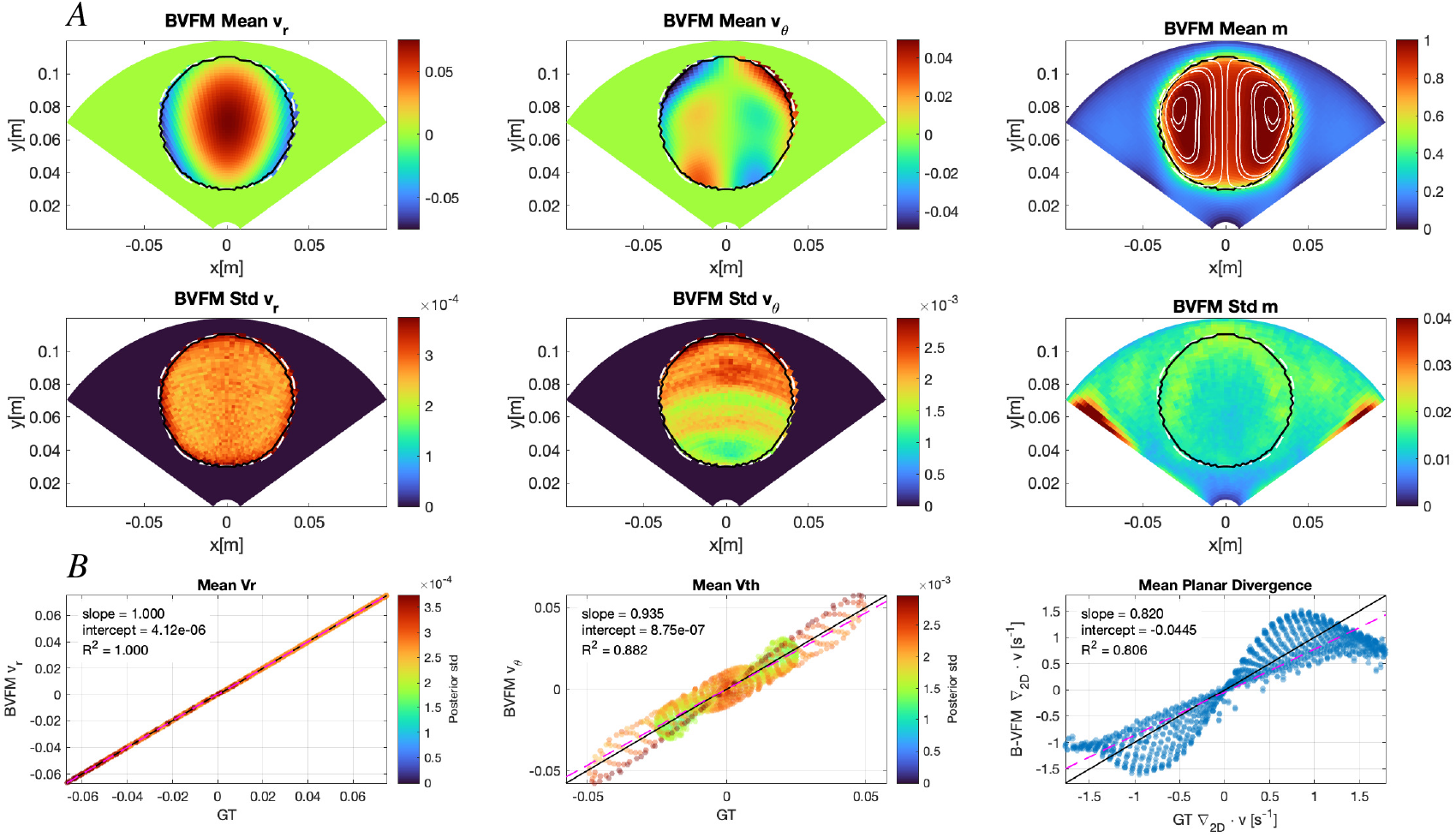
Reconstruction of a Hill’s spherical vortex flow by B-VFM with using mass conservation prior with both learned bias covariance from the HM spherical vortex family. *A*) Spatial distributions of *v*_*r*_, *v*_*θ*_, and the mask *m* with superimposed flow streamlines (left to right). Top row: B-VFM posterior means; bottom row: B-VFM posterior standard deviations. The solid black and white dashed contours denote respectively the boundaries of the reference and inferred mask, defined by *M* = 0.5 and *m* = 0.5 respectively. *B*) Pointwise comparisons of the ground-truth (GT) and inferred velocity components and planar divergence (left: *v*_*r*_, center: *v*_*θ*_; right: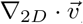). Velocity points are colored according to their posterior standard deviation.——, identity line; 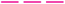, least-squares regression line. Velocity and their posterior standard deviations are given in m,s^*−*1^, divergence has dimensions of s^*−*1^, and the mask is dimensionless.

To independently assess the information learned from data-driven bias and covariance, we also ran B-VFM under a covariance-only data-driven prior, 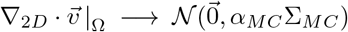, and a bias-only data-driven prior,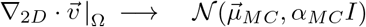. Figure 12*A* presents results from ten independent B-VFM chains using the covariance-only prior. Relative to the original zero-mean, identity-covariance formulation, the learned covariance improved the accuracy of the reconstructed *v*_*θ*_ and kept the inferred mask closer to the reference mask, increasing the Dice score from 0.86 to 0.94. Scatter plots (Figure 12*B*) confirm that the Doppler velocity was recovered exactly (linear slope fit 1.000, *R*^2^ = 1.000), while *v*_*θ*_ was inferred with significantly higher accuracy than in the original formulation (linear slope fit 1.183, *R*^2^ = 0.905). Notably, the planar divergence was reproduced moderately even without prescribing 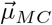. Thus, the spatial covariance structure alone provided useful information about the expected distribution of planar mass-conservation violations. Results from ten independent B-VFM chains using the bias-only prior are shown in Figure SI5 in the Appendix.

**Figure 12:**
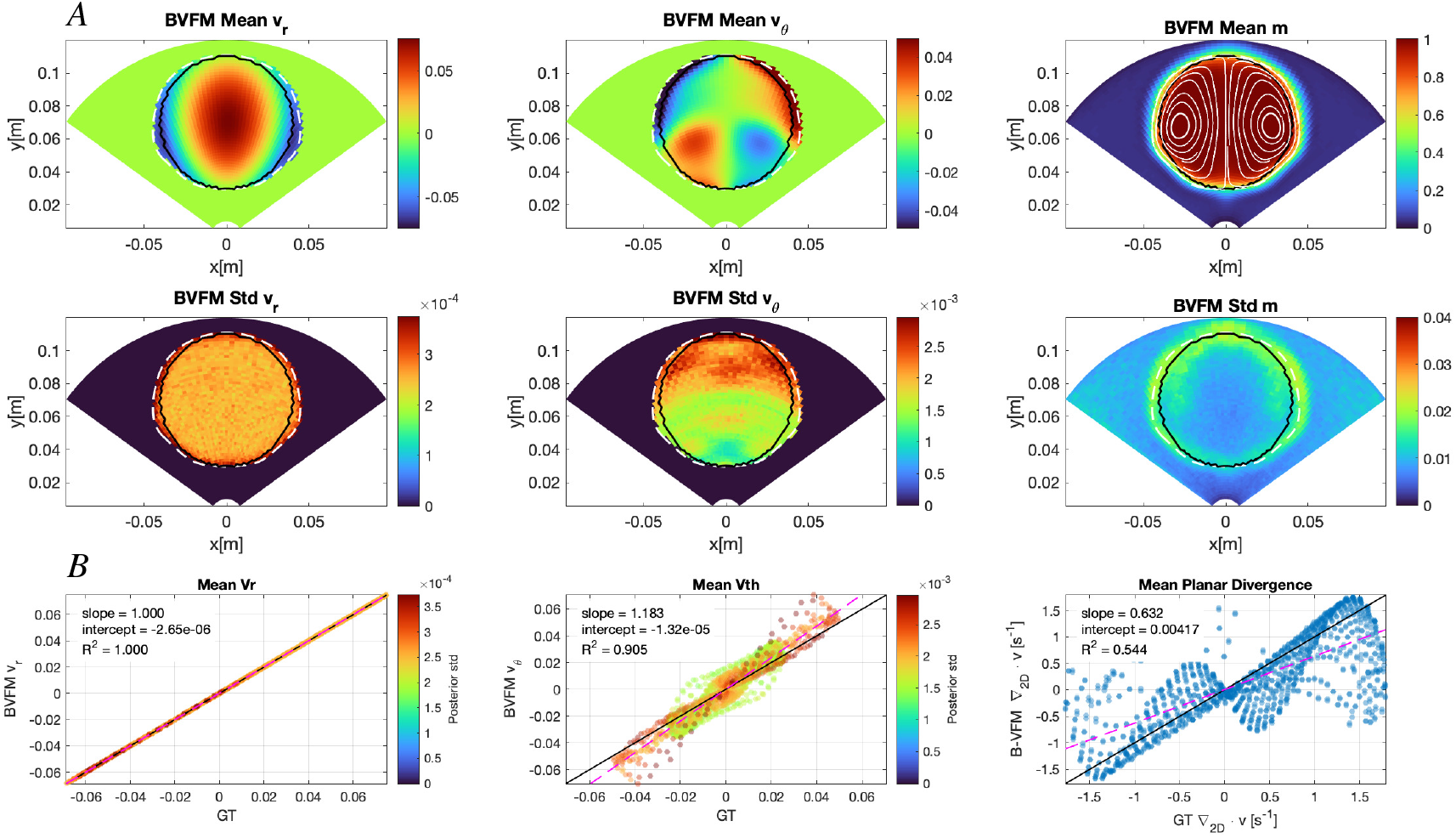
Reconstruction of a Hill’s spherical vortex flow by B-VFM with using mass conservation prior with zero bias and learned covariance from the HM spherical vortex family. *A*) Spatial distributions of *v*_*r*_, *v*_*θ*_, and the mask *m* with superimposed flow streamlines (left to right). Top row: B-VFM posterior means; bottom row: B-VFM posterior standard deviations. The solid black and white dashed contours denote respectively the boundaries of the reference and inferred mask, defined by *M* = 0.5 and *m* = 0.5 respectively. *B*) Pointwise comparisons of the ground-truth (GT) and inferred velocity components and planar divergence (left: *v*_*r*_, center: *v*_*θ*_; right: 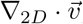). Velocity points are colored according to their posterior standard deviation.——, identity line; 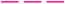, least-squares regression line. Velocity and their posterior standard deviations are given in m,s^*−*1^, divergence has dimensions of s^*−*1^, and the mask is dimensionless.

These trends persisted across the investigated HM family. Figure 13 reports the slopes and coefficients of determination, *R*^2^, from linear fits of reconstructed versus ground-truth *v*_*θ*_, together with Dice scores for the inferred masks, over the range 0 ≤ *κ* ≤ 5. Results are shown for the fully data-driven B–VFM and its covariance-only variant, as well as for the baseline, non-data-driven B–VFM and the reference configurations iVFM[1, 0.1, 10^*−*8^] and iVFM[1, 1000, 10^*−*6^].

**Figure 13:**
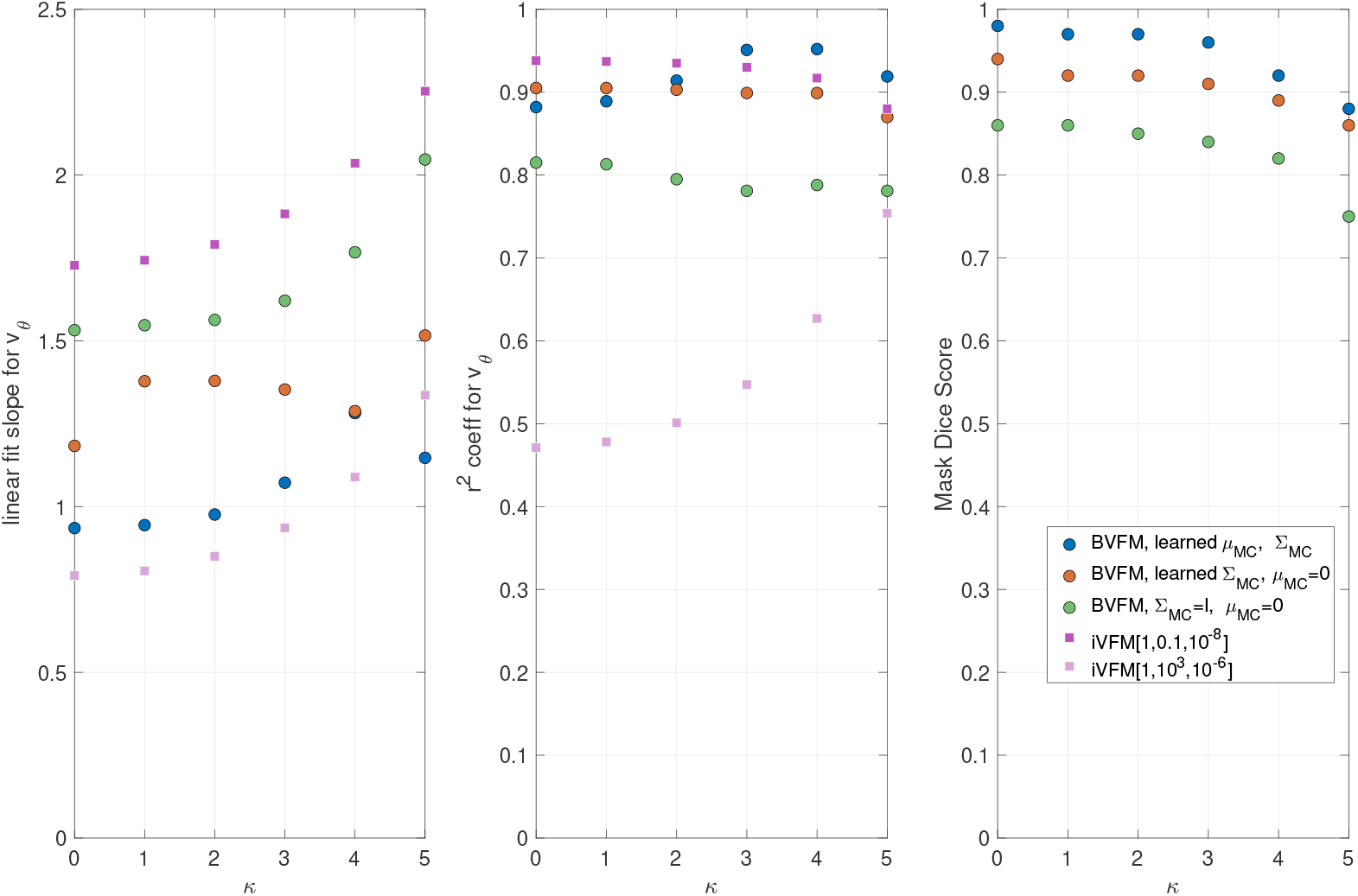
Performance of different BVFM variants across the Hicks–Moffatt (HM) family as a function of the wavenumber parameter *κ*. From left to right, the panels show the slope and coefficient of determination *R*^2^ from linear fits of reconstructed versus ground-truth *v*_*θ*_, and the Dice score of the inferred mask. Results are shown for B–VFM with learned mass-conservation mean and covariance 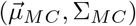, B–VFM with learned covariance only 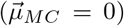, and baseline B–VFM 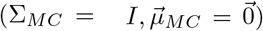. For reference, results from iVFM[1, 0.1, 10^*−*8^] and iVFM[1, 10^3^, 10^*−*6^] are also included. A slope of one indicates no multiplicative bias, whereas larger *R*^2^ and Dice scores indicate better agreement with the ground truth. Dice scores are shown only for the B–VFM variants because iVFM treats the input mask as fixed.

The fully data-driven B–VFM, using the learned 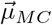 and Σ_*MC*_, performed best overall. Its linear-fit slopes for *v*_*θ*_ remained close to unity, *R*^2^ values were approximately 0.9 across all *κ*, and mask Dice scores were the highest among the methods considered. Covariance-only B–VFM provided the second-best overall performance, although its higher slopes indicated residual overestimation of *v*_*θ*_. Baseline B–VFM and iVFM[1, 0.1, 10^*−*8^] performed similarly and both substantially overestimated *v*_*θ*_; baseline B–VFM produced slopes slightly closer to unity but slightly lower *R*^2^ values compared to iVFM[1, 0.1, 10^*−*8^]. Baseline B–VFM also yielded the lowest mask Dice scores among the B–VFM variants. Finally, iVFM[1, 1000, 10^*−*6^] yielded slopes close to unity but considerable scatter, with *R*^2^ values ranging from approximately 0.5 to 0.75.

## 4 Discussion

Significant progress has been made since the earliest formulations of vector flow mapping (VFM) [22], where the radial velocity (*v*_*r*_) and blood domain (*m*) were fixed to the observed Doppler velocity and segmentation (*V* and *M*), and mass conservation was used to infer the cross-beam velocity (*v*_*θ*_) along each radial arc of the Doppler sector. In these “naïve” implementations, phase unwrapping and spatio-temporal filtering of *V* were performed prior to inference, while *v*_*θ*_ was regularized post hoc using spatial filtering. Since then, VFM has evolved to include least-squares (MAP) formulations with regularization [4, 76], streamfunction–vorticity approaches that promote regularity via a global mass conservation constraint [49, 52], and extensions incorporating physics-informed deep learning [40, 42]. Yet these formulations do not converge to identical solutions, and results remain sensitive to hyperparameters such as smoothing weights, boundary conditions, and segmentation geometry. Moreover, none of these methods utilize flow to correct mask segmentation or propagate uncertainty to the latent variables 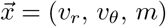.

With VFM reconstructions remaining method- and parameter-dependent, it is becoming important to quantify stability and uncertainty as the technique is extended to increasingly complex secondary analyses [44, 60, 61] and imaging views [1, 3, 16, 28, 30, 39, 50]. Motivated by this need, we formulate B-VFM as a differentiable discrete inverse problem and derive closed-form expressions for its gradient and exact Hessian. The gradient enables efficient MAP estimation, whereas the Hessian reveals how measurement uncertainty, domain geometry, and physical priors govern regularity and couple the latent variables *v*_*r*_, *v*_*θ*_, and *m*. Building on this analysis, we introduce a fully Bayesian implementation of VFM that enables uncertainty quantification, e stablishing a principled foundation for future VFM and multi-modality algorithms, as well as uncertainty-aware secondary analyses. The framework also permits the statistics of model discrepancy to be learned from flow ensembles, allowing to combine physical constraints with empirical information.

### 4.1 Ill-posedness of VFM

Vector flow mapping poses an involved inference problem because the ultrasound cross-beam velocity component, *v*_*θ*_, lacks measurement data. This situation is analogous to the ill-posedness of optical flow in the direction perpendicular to the local image gradient [79], and contrasts with other scenarios, such as regularization of phase-contrast MRI data [81], where there is observational support for all flow components. Several studies have noted that the ill-posed nature of VFM makes it sensitive to implementation details. For instance, Pedrizzetti and Tonti [52] warned that the inferred *v*_*θ*_ could be discontinuous in the absence of global constraints, whereas Assi *et al*. [4] noted that well-posedness requires complete Dirichlet boundary conditions for *v*_*θ*_ on the segmented endocardial boundary (LV mask). However, a formal analysis of VFM regularity had not been presented yet.

Our Hessian analysis makes this ill-posedness explicit. The mass-conservation term has *N*_*r*_ singular modes, each corresponding to an arbitrary additive value of *v*_*θ*_ along a constant-radius arc. Boundary conditions and smoothing act as rank updates to this otherwise singular system: they increase the corresponding eigenvalues and suppress these modes. If neither constraint is effective, the unresolved modes can appear as band-like artifacts in the reconstructed *v*_*θ*_ field, which can be observed in some VFM reconstructions [25, 35].

The analysis also provides precise geometric criteria for regularity. Under free-slip conditions, each constant-*r* arc must intersect the LV boundary at a location with nonzero transverse normal component, *n*_*θ*_. Thus, the free-slip constraint can fail when the mask is open or locally aligned with the *θ* direction on both sides of an arc. Such configurations are more likely in the presence of segmentation notches [25], intracardiac shunts, or in chambers such as the RV, where inflow and outflow tracts can align with the *θ* direction [35]. Under no-slip conditions, regularity instead requires that each arc contain at least one segmented endocardial pixel. This result suggests that no-slip conditions are more robust to local mask geometry, despite being less consistent with the mass-conservation model.

The Laplacian smoothing operator suppresses the arcwise modes left undetermined by transverse mass conservation; the only shared null mode is the globally constant mode, which is constrained by the boundary conditions. This conclusion, however, does not automatically carry over to all discrete operators. For example, constructing the radial Laplacian by applying a centered first-difference twice yields a wider stencil that decouples even and odd radial mesh points. The resulting checkerboard mode belongs to the null spaces of both the discrete smoothing and transverse mass-conservation operators. It therefore remains unpenalized by smoothing and can appear as alternating radial bands in *v*_*θ*_, a phenomenon we observed in our numerical experiments and which is closely related to the even–odd decoupling artifact encountered in computational fluid dynamics [56]. Of note, the null-space argument also clarifies why artifact suppression requires smoothing to be included as a prior during inference. A filter applied after reconstructing *v*_*θ*_ may reduce visual noise, but it cannot regularize the inverse problem or remove its null modes.

### 4.2 Hierarchical Bayesian VFM

Although the Navier–Stokes equations include a viscous term that promotes smoothness, the smoothing typically applied in VFM is not a direct physical constraint, leaving ambiguity over how much relative weight should be assigned to the different constraints. The L-curve is a popular heuristic for tuning regularization parameters [26] that was previously adopted in least-squares VFM [4]. While simple and widely used, its corner is often ambiguous, it is sensitive to noise and scaling, and multidimensional extensions are computationally demanding and difficult to interpret [7]. Most importantly, the L-curve provides only point estimates of the weights, without a framework for uncertainty quantification. Moreover, scalar weights cannot naturally encode spatially varying confidence or correlations within individual constraints, nor quantify interactions among constraints. In contrast, a hierarchical Bayesian approach can prescribe covariance structures that represent those details while treating their overall scales as hyperparameters estimated jointly with the latent vector 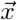 [77].

Motivated by these considerations, we adopted a hierarchical Bayesian formulation in which both 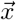 and 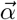 were treated as random variables. To emulate their joint posterior, we used Markov Chain Monte Carlo (MCMC), sequentially updating 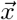 and 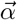 by sampling from their conditional distributions [23]. MCMC provides a principled framework for exploring high-dimensional posteriors and quantifying uncertainty in both the physical model and the data. In B-VFM, however, the conditional posteriors are challenging because the segmentation mask *m* is latent, introducing nonlinearities that render the priors non-Gaussian. The boundary-condition prior 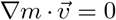 illustrates this difficulty, as it tightly couples the mask and the flow field. To approximate such conditionals by Gaussian distributions, we employed local Laplace expansions [73] around the MAP estimate of the conditional log-posterior, 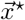, with precision matrices 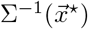 obtained from the log-posterior Hessian. Similar Laplace-based approximations have been applied in medical imaging, including nonrigid registration [78], quantitative MRI multi-parameter mapping [6], and radiation image reconstruction [38], demonstrating their value for tractable uncertainty quantification in high-dimensional inverse problems.

Analytical expressions for the residual Jacobians enabled us to construct a Gauss–Newton approximation of 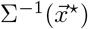 and evaluate its action through Jacobian–vector products. Samples from the resulting Laplace approximation could therefore be generated using iterative linear solves, without explicitly assembling or inverting the full precision matrix, substantially reducing computational cost and improving numerical stability. This is important because numerical stabilization of operations involving the precision matrix often increases its diagonal dominance, which can attenuate the off-diagonal mask–velocity coupling that enables flow-informed segmentation correction. We observed this issue in our numerical experiments when using an incomplete Cholesky factorization of the precision matrix.

For hyperparameter sampling, we used Gamma-distributed hyperpriors, which are appealing because their conjugacy ensures that the conditional hyperparameter posteriors remain Gamma distributed and can be sampled directly. We selected shape and rate parameters so that these hyperpriors were effectively non-informative over the relevant hyperparameter ranges. The substantial Kullback– Leibler divergences between the hyperpriors and posterior hyperparameter distributions indicated that the observations and physical constraints provided considerable information in determining the inferred weights. Overall, the Gibbs sampler exhibited rapid and robust convergence, consistent with the shrinkage properties of Gamma hyperpriors in high-dimensional latent spaces 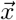 [23].

### 4.3 Flow Informed Segmentation Refinement

Traditional VFM formulations [4, 21, 22, 33, 49, 69, 76] treat the LV mask as fixed, effectively imposing *m* = *M* as a deterministic constraint. If the same assumption were applied to B-VFM, the conditional structure would simplify substantially: the conditional posterior for the velocity field would be Gaussian, and the hyperparameters could then be updated directly by Gibbs sampling without MAP estimation. Even if a MAP estimate were computed, it would reduce to solving a linear system, as in least-squares VFM [4, 76], dramatically accelerating the computations.

Such simplification, however, would remove the possibility of using velocity data to refine or correct blood pool segmentation, an idea recently explored in different forms [15, 37]. In the LV, this trade-off is especially relevant in pathologies where delineation of the endocardial border is challenging or unreliable. In dilated cardiomyopathy, the myocardium is thinned, reducing contrast with the blood pool and producing poorly defined endocardial borders in the apical view typically used for VFM [53]. In LV non-compaction, prominent trabeculations and deep recesses obscure the boundary, and color Doppler is often leveraged to improve diagnostic accuracy [67]. In patients with apical hypertrophic cardiomyopathy or apical aneurysms, the apex may nearly or completely obliterate during systole, with segmentation further complicated by asymmetric wall thickening, aberrant trabeculation, and displaced papillary muscles [43].

In B-VFM, the boundary-condition operator depends on *m*, mass conservation is enforced only inside *m*, and the Doppler misfit is evaluated within *m*. The most direct coupling between flow and segmentation arises from the boundary-condition residual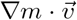. Because *∇m* is normal to the level sets of the mask, minimizing this residual favors an inferred boundary whose tangent is aligned with the local flow. The residual can therefore be reduced by adjusting either the velocity field or the location and orientation of the boundary, allowing velocity information to add or remove pixels and reshape the inferred mask. This two-way interaction is explicitly encoded in the off-diagonal velocity–mask blocks of the precision matrix.

Our benchmarks illustrate both the value and the potential limitations of this coupling. B-VFM corrected the artificial defect in the Lamb–Chaplygin mask, demonstrating that flow information can improve segmentation when the physical model is consistent with the data. Conversely, when the observations cannot be reconciled with the physical priors, part of the model discrepancy may be absorbed through deformation of the inferred mask. This behavior was evident in the 3D Hicks-Moffatt benchmark, for which the planar mass-conservation cannot be achieved without violating the boundary conditions, to which BVFM responded by expanding the inferred mask. Although the segmentation likelihood and mask-smoothing prior provide anchoring, they may not be sufficient under substantial model mismatch. We therefore introduced optional hard interior and exterior constraints that fix pixels confidently identified as blood or tissue while allowing the boundary to adapt in the rest of the image. Our numerical experiments showed that these constraints substantially mitigated spurious mask deformation and often produced more meaningful posterior uncertainty maps.

### 4.4 Data-informed modeling of physical-model discrepancy

Similar to previous VFM methods [4, 76], the baseline B-VFM formulation assigns the planar mass-conservation residual a zero mean and a covariance proportional to the identity matrix. This corresponds to a white-noise prior that treats residuals at different locations as spatially independent and equally variable. Because this prior contains no information about coherent divergence patterns produced by out-of-plane transport, it biases the reconstruction toward 2D incompressibility. Our Hicks– Moffatt benchmarks showed that this model mismatch can substantially overestimate the transverse velocity.

A more general formulation can incorporate the mean and covariance of the planar-divergence residual learned from 3D velocity data, such as 4D flow MRI. The learned covariance relaxes the planar incompressibility constraint in a spatially structured manner, potentially improving both reconstruction accuracy and uncertainty quantification. Because the empirical covariance is represented as a diagonal matrix plus a low-rank ensemble contribution, its inverse is applied without storing or factorizing a full spatial covariance matrix, thus conserving computational efficiency.

Our HM results show that incorporating both the learned mean and covariance produced the most accurate reconstruction, while the covariance-only prior captured much of this improvement. The latter may be particularly advantageous in clinical applications, where interpatient variability and reduced spatial coherence of the velocity fields may cause patient-specific divergence patterns to cancel in the population mean. In that setting, prescribing a fixed mean could be overly restrictive. The present results, however, only provide an in-family proof of principle for the synthetic Hicks–Moffatt family of vortex dipoles. Establishing generalization will require independent training and evaluation ensembles and, ultimately, priors learned from experimental, computational, or clinical 3D flows.

### 4.5 Extensions of B-VFM

The hierarchical Bayesian framework introduced here is inherently flexible, and its computational cost is relatively insensitive to the number of hyperparameters, making it suitable for customization with additional priors and observation modalities. The formulation can be extended to 3D settings [75], where coupling across slices or volumetric Doppler acquisitions would be handled within the same probabilistic framework. It could also accommodate multimodal fusion: data and/or uncertainty estimates from echo-PIV [66] or 4D flow MRI [19] could be incorporated as additional observations.

On the computational side, operator-discretization iterative linearization (ODIL) [36] and physics-informed neural networks (PINNs) [55] provide alternative ways of formulating PDE-constrained inference and could be exploited to facilitate maximum a posteriori (MAP) estimation in 3D and time-resolved B-VFM. Dimensionality-reduction strategies, from Karhunen–Loève expansions [46] to likelihood-informed parameter subspaces (LIPS) [17], have also been developed to approximate high-dimensional posteriors in more tractable subspaces. Their effectiveness depends critically on knowledge of the precision matrix to identify data-informed directions, and the availability of an analytic expressions for the precision matrix in B-VFM makes such reductions especially well-suited for scalable posterior approximation.

## 5 Conclusion

We introduced B–VFM, a flexible hierarchical Bayesian framework that jointly reconstructs LV velocity fields, refines blood-pool segmentation, and quantifies uncertainty from color-Doppler data. Its discretized formulation is differentiable with closed-form gradient and Hessian operators, providing a reusable foundation for optimization, uncertainty propagation, regularity analysis, data fusion, and future higher-dimensional extensions. The framework further allows systematic physical-model discrepancies to be learned from flow ensembles through structured prior means and covariances, as demonstrated here using synthetic 3D vortices. Future work will validate B–VFM using clinical data and evaluate it with more complex benchmark flows.

## Acknowledgments

This work was supported by the National Science Foundation Graduate Research Fellowship under grant No.DGE-2140004 (to CN), AHA Collaborative Award 25CSA1421482 (to JCdA), and by the Spanish Research Agency (AEI, grant number PID2023-146861OB-I00, to PML). JCdA would like to acknowledge support from the James B. Morrison endowment. We appreciate fruitful discussions with Oscar Flores about phase unwrapping.

## Supporting information

### SI1 The B-VFM loss and gradient

The MAP step of the sampler described in Section 2.2.1 minimizes the conditional negative log-posterior with respect to the latent state 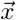. Here, we express this loss in terms of the residuals associated with the likelihood and prior models and derive its gradient.

Let *N* = *N*_*r*_*N*_*θ*_, use *δ*(*a*) = diag(*a*), and write the discrete divergence operator as

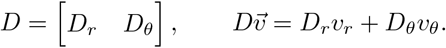

The radial and azimuthal components of the discrete gradient operator are denoted by *G*_*r*_ and *G*_*θ*_.

For each likelihood or prior term, we separate the scalar precision *α*_*i*_ from any prescribed spatial covariance structure by writing

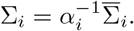

In the expressions below, 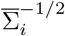 denotes a factor satisfying

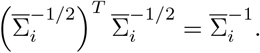

For spatially white models 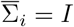.

For fixed hyperparameters, the conditional negative log-posterior is

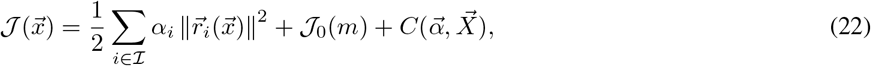

where

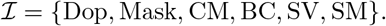

The term *J*_0_(*m*) contains normalization factors that depend on the latent mask and is specified below. The remaining Gaussian normalization terms are collected in 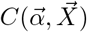, which is independent of 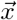 for fixed 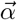 Thus, *C* does not affect the MAP estimate, although its 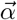-dependent terms are retained when updating the hyperparameters.

We denote the unweighted Doppler, mask, mass-conservation, and boundary-condition mismatches by

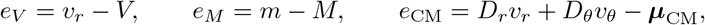

and

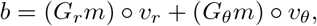

where ° denotes elementwise multiplication. In the baseline mass-conservation prior, ***µ***_CM_ = 0 and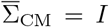. In the data-driven prior, ***µ***_CM_ is the ensemble-mean divergence and 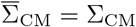 is the regularized covariance constructed from the ensemble matrix *A* in Section 2.3.1.

The six residual vectors are

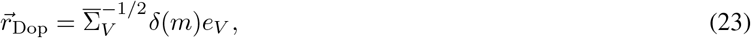

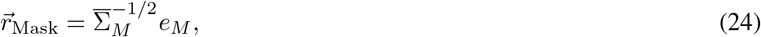

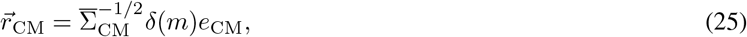

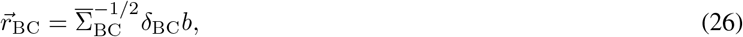

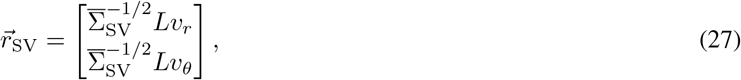

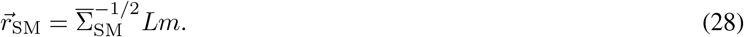

Here, *δ*_BC_ is the fixed diagonal operator that suppresses boundary-condition enforcement near open valves or at other prescribed locations.

The factors |*δ*(*m*)| in the mass-conservation prior and Doppler likelihood contribute to the Jacobian as

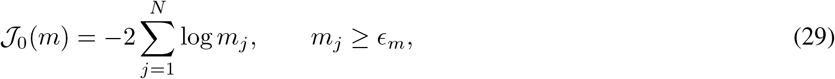

where *ϵ*_*m*_ = 10^*−*6^ is a hard lower bound on the mask. Therefore,

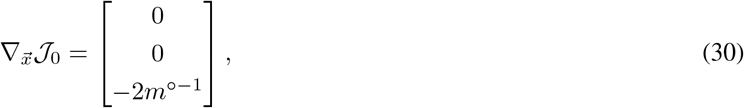

where *m*^*°−*1^ denotes the elementwise reciprocal of *m*.

For each residual, let

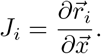

Differentiating equations (23)–(28) gives

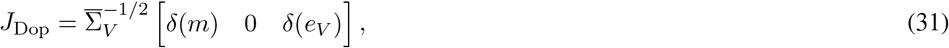

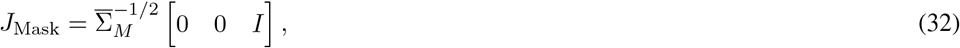

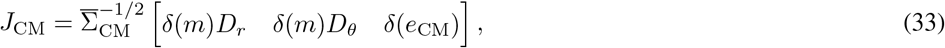

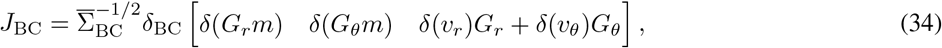

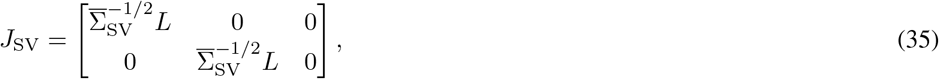

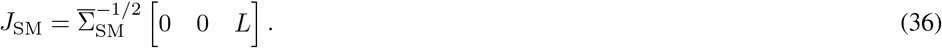

The gradient of each weighted squared residual is

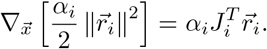

Consequently, the gradient used for MAP optimization is

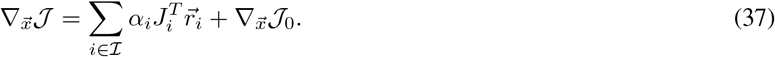

The residuals and Jacobians in equations (23)–(36) are evaluated at the current iterate during MAP optimization.

### SI2 Exact Hessian, Gauss–Newton approximation, and Neumann-series approximation to its inverse

The Laplace approximation requires the local curvature of the conditional negative log-posterior at the MAP estimate. Here, we derive the exact Hessian of the B-VFM loss and compare it with the Gauss–Newton Hessian used for posterior sampling. At the conditional MAP estimate 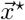, the exact Hessian of the negative log-posterior is

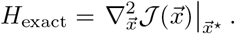

For each squared residual in equation (22),

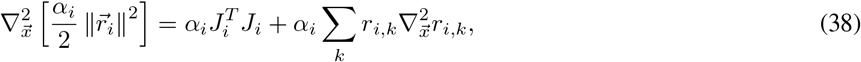

which has a part that can be absorbed in the Gauss-Newton approximation and a part that cannot. Specifically, the Gauss– Newton approximation retains the first term in equation (38) and the exact Hessian of *J*_0_:

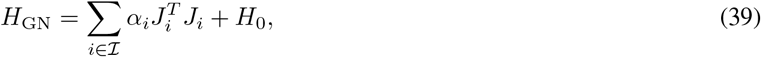

where

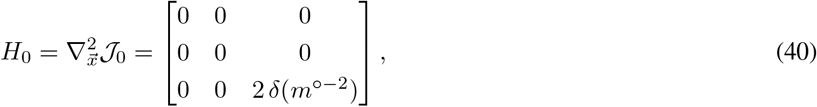

and *m*^*°−*2^ denotes the elementwise inverse square of *m*.

0 0 2 *δ*(*m*^*°−*2^)

The mask, velocity-smoothing, and mask-smoothing residuals are linear in 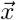, so their second derivatives vanish. Therefore, their Gauss-Newton approximation is exact. The exact Hessian is therefore

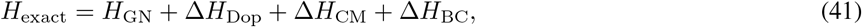

where the Δ*H* terms represent the contributions from the Doppler, mass-conservation, and boundary-condition residuals are nonlinear because they couple the velocity and mask. For these terms, we define

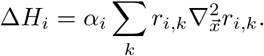

Under the block ordering 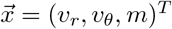, the only nonzero block of the Doppler correction is

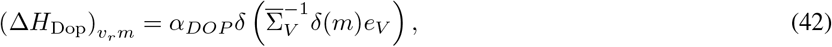

together with its transpose 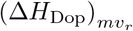.

The nonzero upper-triangular blocks of the mass-conservation correction are

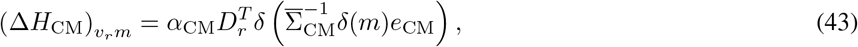

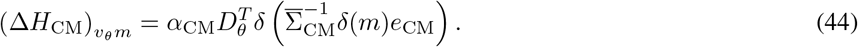

The corresponding lower-triangular blocks are obtained by transposition.

Similarly, the nonzero upper-triangular blocks of the boundary-condition correction are

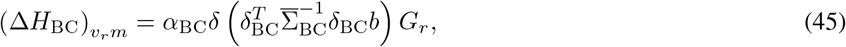

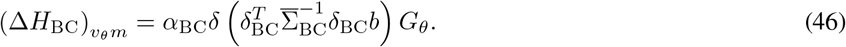

Again, the corresponding lower-triangular blocks are obtained by transposition. All unlisted blocks of the three correction matrices are zero.

Equivalently, the complete correction can be displayed in block form as

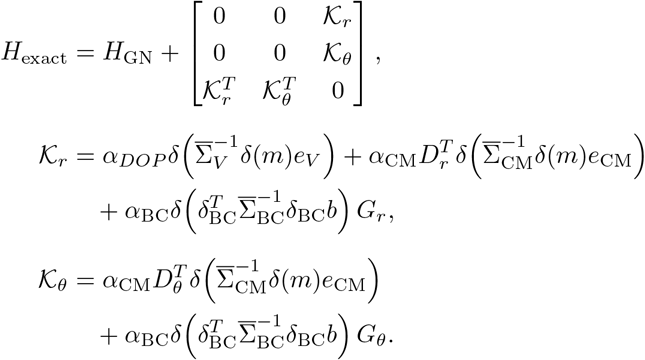

The correction matrices are proportional to the corresponding residuals and vanish when those residuals are zero. They are not guaranteed to be positive semidefinite. B-VFM therefore uses *H*_GN_ in the Laplace approximation, giving

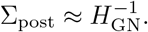

When hard mask constraints are imposed, the Hessian and covariance expressions are restricted to the unconstrained coordinates.

Next, we derive a block-wise Neumann-series approximation to *H*^*−*1^ that is useful to analyze the ill-posedness of VFM. Because *v*_*r*_ and *m* are directly observed whereas *v*_*θ*_ is not, the observation terms dominate the (*r, r*) and (*m, m*) Hessian blocks when the Doppler and mask errors are small. Let *σ* denote the common order of magnitude of these observation covariances, such that *σ*^2^ *∼* Σ_*V*_ *∼* Σ_*M*_ *≪* 1. Representing the overall strength of the VFM priors by a generic precision *α*, the leading-order Hessian has the block form

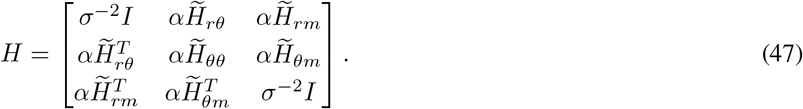

Here, the matrices 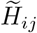 have been normalized to have norms comparable to that of the identity matrix. Contributions of the priors to the (*r, r*) and (*m, m*) blocks are smaller than the corresponding observation terms by a factor of order *ασ*^2^ and have therefore been omitted from Equation (47).

We decompose the Hessian as

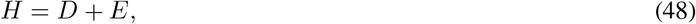

where

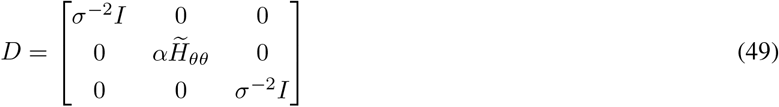

contains the leading diagonal blocks, and

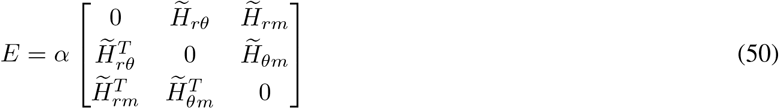

contains the cross-variable couplings. Provided that *D* is nonsingular, the Hessian can be factored as

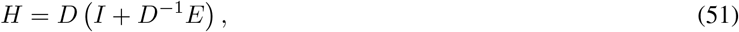

and therefore

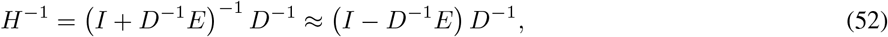

keeping the first-order term of the Neumann series approximation. Since

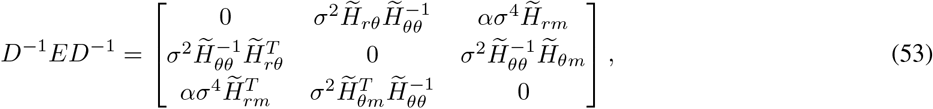

then

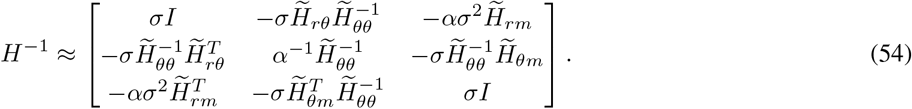

### SI3 Phase Unwrapping Can Be Decoupled from Inference in B-VFM

The color Doppler maps *V* (*r, θ*) used as input to B-VFM may exhibit aliasing artifacts due to phase wrapping of the Doppler phase when the true radial velocity exceeds the encoding limit *V*_enc_. These artifacts create ambiguity in the observed phase, Φ = *πV/V*_enc_. In pixels affected by aliasing, the true (*φ*) and observed phase differ by an integer multiple of 2*π*:

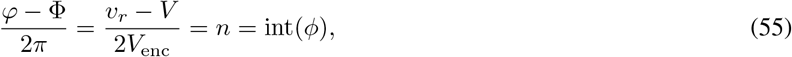

where *n* ∈ ℤ is the aliasing index, and *ϕ* ∈ ℝ is its real-valued extension.

To account for this ambiguity within the Bayesian framework, we introduce *ϕ* as an additional latent variable in the forward model 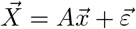, modifying the observation equation to:

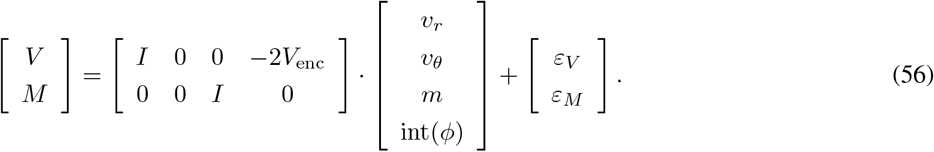

When *n* = *ϕ* = 0, there is no phase wrapping, and the model reduces to the scenario described in the Main Text.

To correct phase-wrapping artifacts, we incorporate prior knowledge on the spatial structure of *ϕ* following the approach of Loecher et al. [41]:

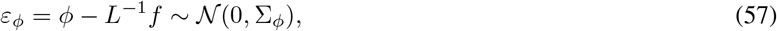

where *L* is the Laplace operator, and

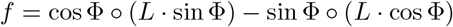

is a nonlinear function of Φ, itself a function of *V*. The prior covariance 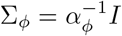 is diagonal, with precision hyperparameter *α*_*ϕ*_. The corresponding probability density function is:

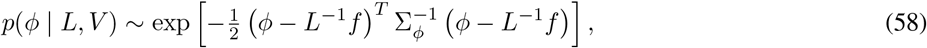

introducing a multiplicative term in the conditional posterior (eq. 10 in the Main Text). In addition to this prior, the likelihood for *V* is modified to:

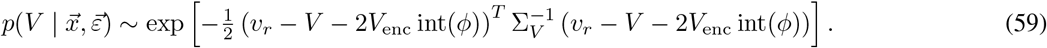

Differentiating the negative log conditional posterior with respect to *ϕ* yields the following MAP condition:

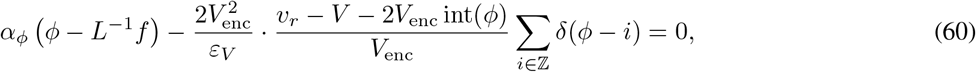

where the final term is a Dirac comb used to approximate the distributional derivative of int(*ϕ*) with respect to *ϕ*. This derivative is zero almost everywhere (for 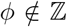), effectively decoupling the MAP estimate of *ϕ* from other latent variables and yielding *ϕ* = *L*^*−*1^*f*. When *n* = *ϕ* ∈ ℤ, equation 60 reduces to the aliasing identity 2*V*_enc_*n* = *v*_*r*_ − *V*. These results hold independently of *α*_*ϕ*_ and shows that phase unwrapping can be performed as a standalone preprocessing step prior to executing the B-VFM algorithm while retaining Bayesian rigor.

### SI4 Supplementary Figures

**Figure SI1:**
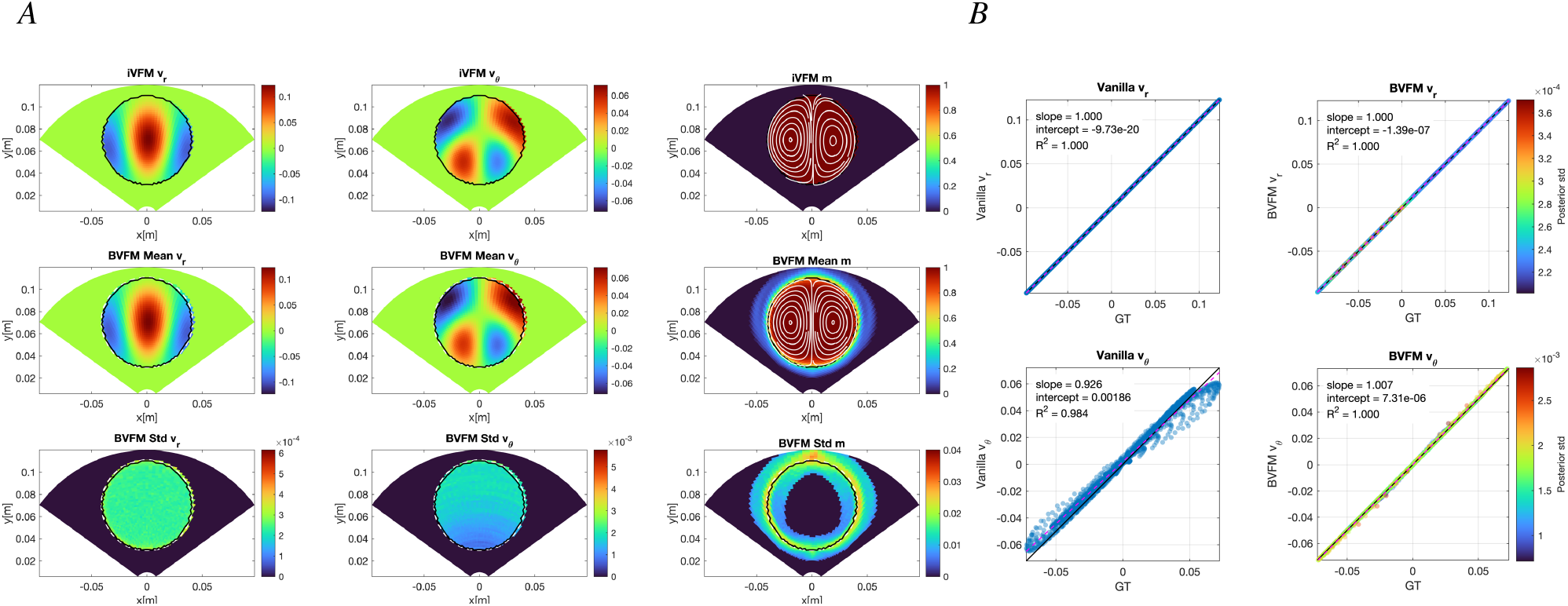
Reconstruction of the ideal, noiseless LC dipole with hard mask constraints. *A*) Spatial distributions of *v*_*r*_, *v*_*θ*_, and the mask *m* with superimposed flow streamlines (left to right). Top row: iVFM [1, 0.1, 10^*−*6^] reconstruction and reference binary mask; middle row: B-VFM posterior means; bottom row: B-VFM posterior standard deviations. The solid black and white dashed contours denote respectively the boundaries of the reference and inferred mask, defined by *M* = 0.5 and *m* = 0.5 respectively. *B*) Pointwise comparisons of the ground-truth (GT) and reconstructed velocity components (top: *v*_*r*_, bottom: *v*_*θ*_; left: iVFM[1,1,10^*−*6^], right: B-VFM). B-VFM points are colored according to their posterior standard deviation. ——, identity line; 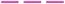, least-squares regression line. Velocity and their posterior standard deviations are given in m,s^*−*1^, and the mask is dimensionless.

**Figure SI2:**
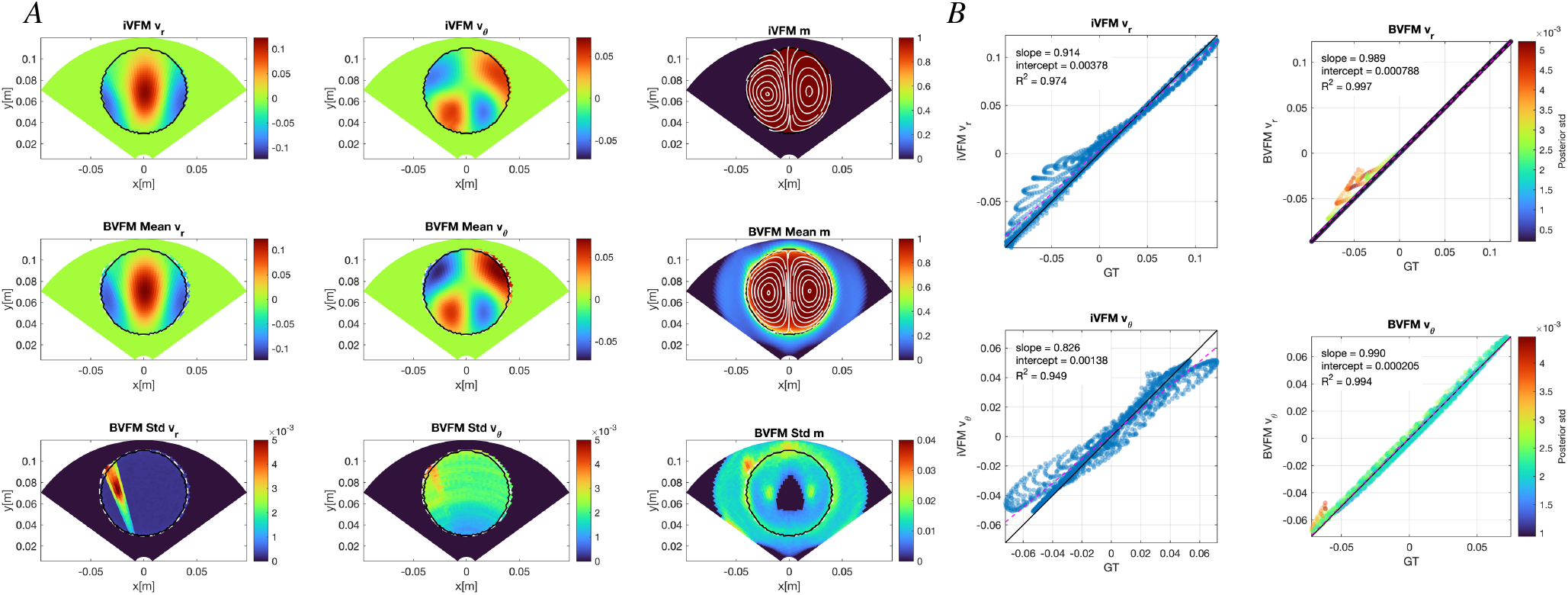
Reconstruction of a LC dipole with a twinkling Doppler artifact (see Figure 7*A*) with hard mask constraints. *A*) Spatial distributions of *v*_*r*_, *v*_*θ*_, and the mask *m* with superimposed flow streamlines (left to right). Top row: iVFM [1, 0.1, 10^*−*5^] reconstruction and reference binary mask; middle row: B-VFM posterior means; bottom row: B-VFM posterior standard deviations. The solid black and white dashed contours denote respectively the boundaries of the reference and inferred mask, defined by *M* = 0.5 and *m* = 0.5 respectively. *B*) Pointwise comparisons of the ground-truth (GT) and reconstructed velocity components (top: *v*_*r*_, bottom: *v*_*θ*_; left: iVFM[1,0.1,10^*−*4^], right: B-VFM). B-VFM points are colored according to their posterior standard deviation. ——, identity line; 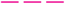, least-squares regression line. Velocity and their posterior standard deviations are given in m,s^*−*1^, and the mask is dimensionless.

**Figure SI3:**
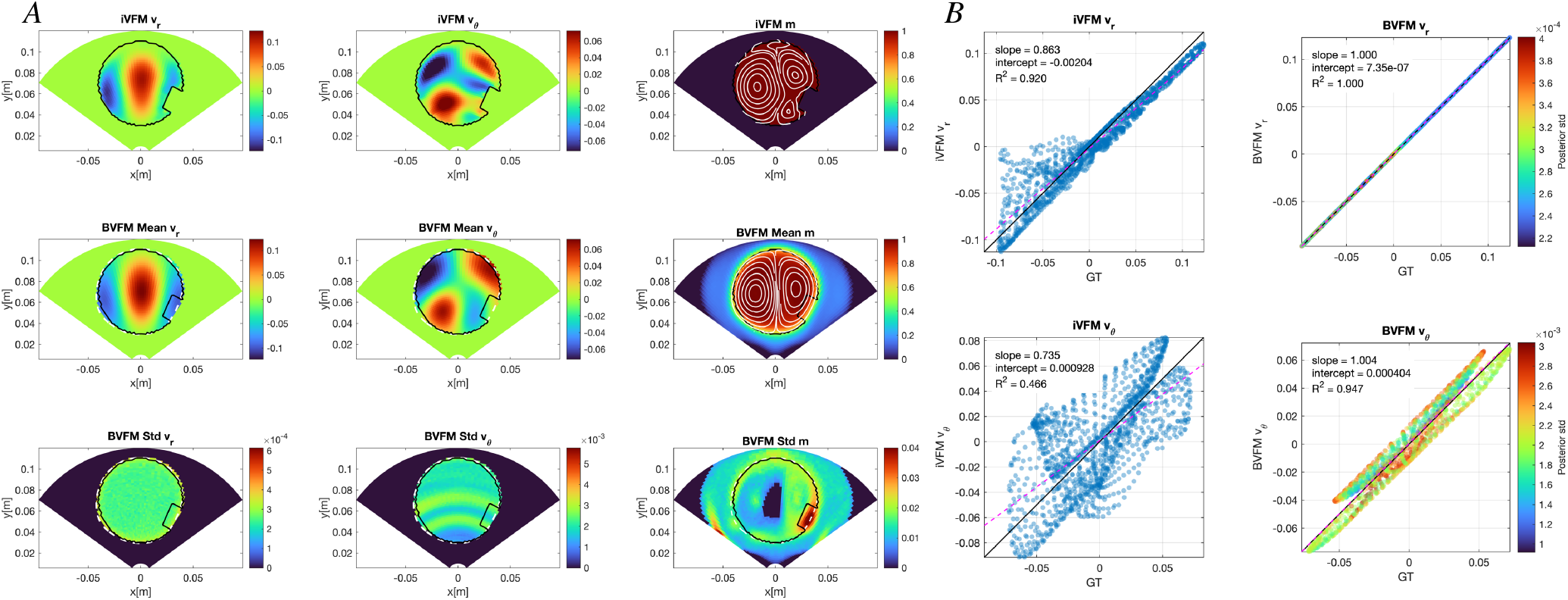
Reconstruction of a Lamb–Chaplygin (LC) dipole with a twinkling Doppler artifact with hard mask constraints. *A*) Spatial distributions of *v*_*r*_, *v*_*θ*_, and the mask *m* with superimposed flow streamlines (left to right). Top row: iVFM [1, 10^4^, 10^*−*6^] reconstruction and reference binary mask; middle row: B-VFM posterior means; bottom row: B-VFM posterior standard deviations. The solid black and white dashed contours denote respectively the boundaries of the reference and inferred mask, defined by *M* = 0.5 and *m* = 0.5 respectively. *B*) Pointwise comparisons of the ground-truth (GT) and reconstructed velocity components (top: *v*_*r*_, bottom: *v*_*θ*_; left: iVFM [1, 10^4^, 10^*−*6^], right: B-VFM). B-VFM points are colored according to their posterior standard deviation.—— identity line; 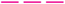, least-squares regression line. Velocity and their posterior standard deviations are given in m,s^*−*1^, and the mask is dimensionless.

**Figure SI4:**
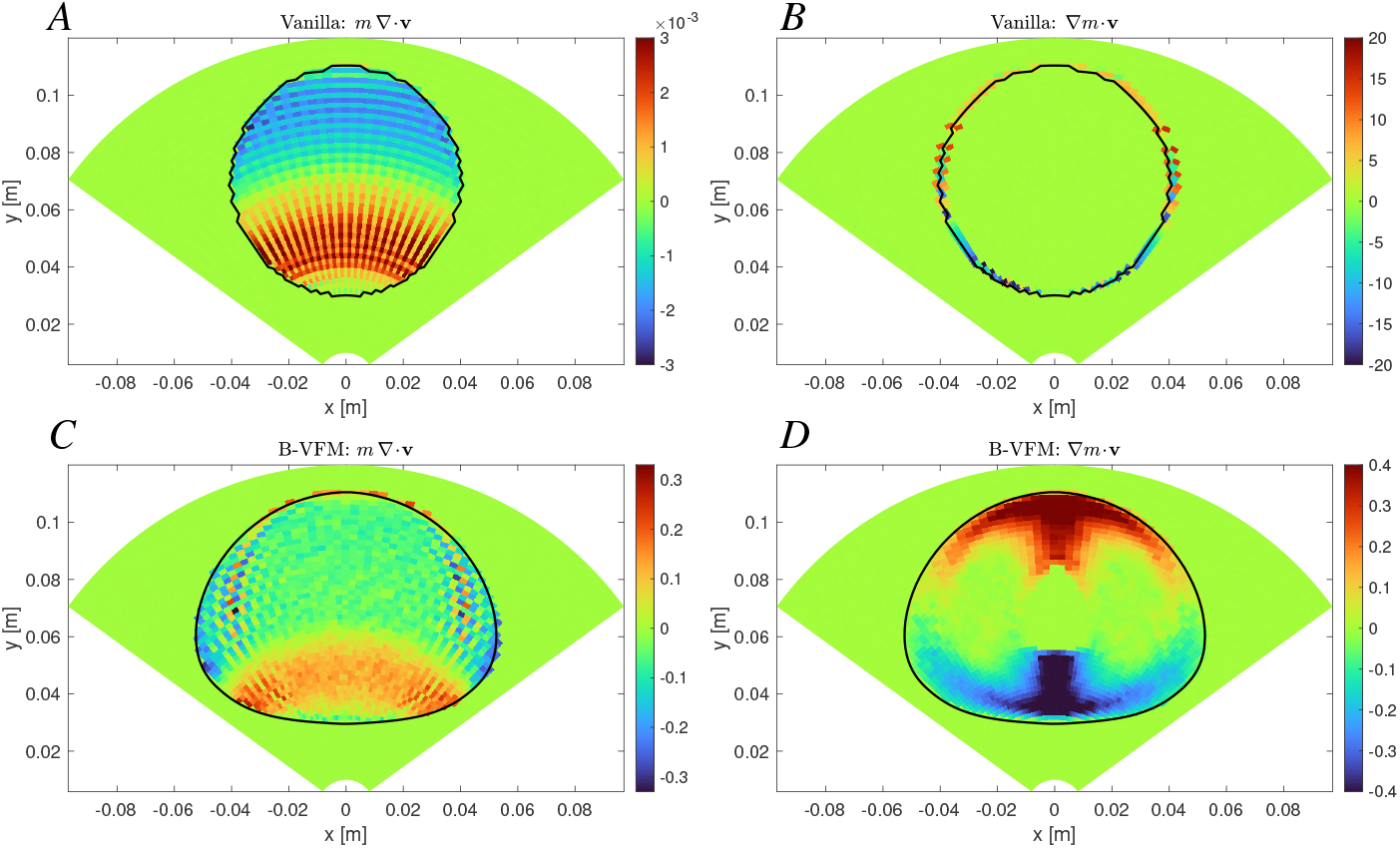
Residuals of 2D mass-conservation and boundary conditions constraints for the Hill’s vortex VFM reconstruction. *A*) 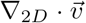, iVFM [1, 0.1, 10^*−*4^] (labeled “Vanilla”), units: 1/s. *B*)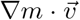, iVFM [1, 0.1, 10^*−*4^], units: 1/s. *C*)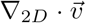, B-VFM, units: 1/s. *D*) 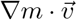, B-VFM, units: 1/s.

**Figure SI5:**
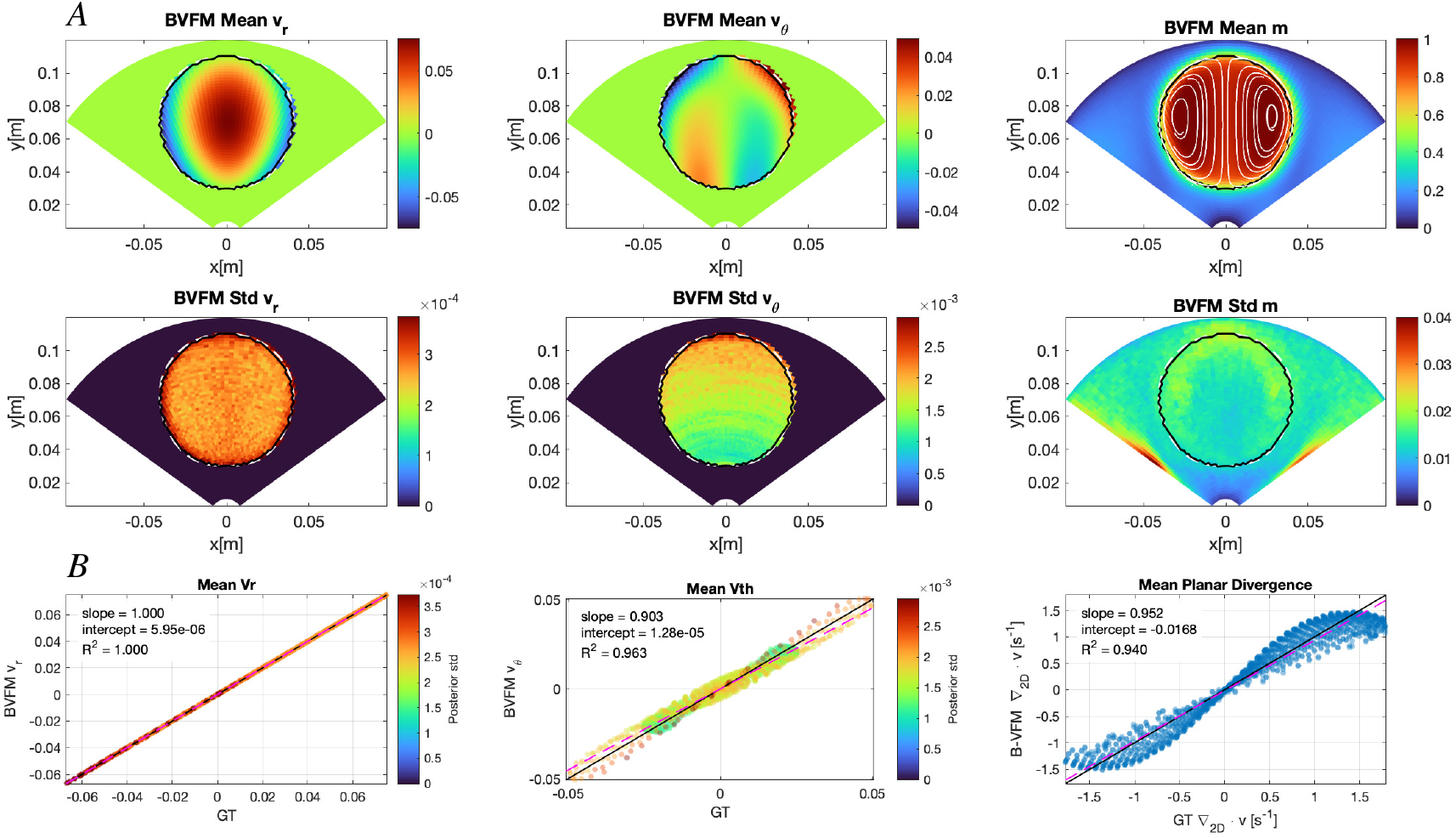
Reconstruction of a Hill’s spherical vortex flow by B-VFM with using mass conservation prior with white noise covariance and learned bias from the HM spherical vortex family. *A*) Spatial distributions of *v*_*r*_, *v*_*θ*_, and the mask *m* with superimposed flow streamlines (left to right). Top row: B-VFM posterior means; bottom row: B-VFM posterior standard deviations. The solid black and white dashed contours denote respectively the boundaries of the reference and inferred mask, defined by *M* = 0.5 and *m* = 0.5 respectively. *B*) Pointwise comparisons of the ground-truth (GT) and inferred velocity components and planar divergence (left: *v*_*r*_, center: *v*_*θ*_; right: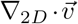). Velocity points are colored according to their posterior standard deviation.———, identity line; 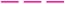, least-squares regression line. Velocity and their posterior standard deviations are given in m,s^*−*1^, divergence has dimensions of s^*−*1^, and the mask is dimensionless.

